# Noninvasive Focal Gene Delivery of Functional Neural Actuators to the Primate Spinal Cord using Focused Ultrasound

**DOI:** 10.64898/2026.09.05.749619

**Authors:** M. R. Corigliano, S. S. Guretse, N. T. Nosenchuck, I. Zimmermann Rollin, L. Letica, D. Szczupak, V. P. Campos, L. Liang, M.C. Noh, T. K. Hitchens, E. Pirondini, R. Seal, M. K. Lin, D.J. Schaeffer

## Abstract

Pathologies of the spinal cord – from degenerative diseases to chronic pain – represent a substantial global health burden. Although surgical and pharmacological treatments for spinal cord pathologies have advanced considerably, therapies capable of addressing the cellular mechanisms underlying these conditions remain limited. Viral gene therapies present a compelling alternative, allowing for delivery of therapeutic genes that directly target the pathological processes in specific cell-types. Effective viral delivery to the spinal cord, however, remains constrained by a fundamental tradeoff between procedural invasiveness and spatial precision. As such, there remains a need for clinically tractable methods – that are both noninvasive and focal – to deliver gene therapeutics across the restrictive vascular boundaries of the blood-spinal cord barrier (BSCB). Here, we demonstrate noninvasive, focal disruption of the BSCB and delivery of systemically administered chemogenetic gene payloads to the cervical and thoracic spinal cord in marmoset nonhuman primates (*Callithrix jacchus*) using focused ultrasound (FUS). Through systematic testing of ultrasonic pressures and central frequencies, we establish a FUS parameter set optimized for robust, spatially constrained molecular delivery across the marmoset BSCB. Using these optimized parameters, we show that FUS BSCB disruption permitted focal penetrance of systemically administered viral vectors for transduction of both fluorescent transgenes and excitatory chemogenetics within targeted spinal segments. Positron emission tomography (PET) imaging following chemogenetic actuation revealed significantly increased metabolic demand within the targeted region of the spinal cord, demonstrating *in vivo* evidence of functional transgene expression. Behavioral and histopathological assessments demonstrated preserved neurological function and tissue integrity, supporting the safety of FUS BSCB disruption and viral delivery in nonhuman primates. To facilitate broad application of this platform for noninvasive delivery of receptor-based gene therapeutics in marmosets, we generated an ultra-high-resolution (74 µm) multimodal MRI/CT spinal cord template for precise targeting and anatomical localization. We also provide open-access engineering drawings and CAD files for our M-FRAME system (Marmoset Fixation and Reorientation Apparatus for Multimodal Experiments), enabling precise and repeatable spinal targeting without surgical fixation. Together, these results establish FUS-mediated BSCB disruption as a safe and effective approach for noninvasive, focal gene delivery to the primate spinal cord.

## Introduction

Pathologies of the spinal cord, including spinal cord injury and pain-related disorders, represent a significant global health burden^1–5^. Despite major advances in surgical and pharmacological treatments for spinal cord pathologies, therapeutic options capable of addressing the cellular etiology of these life altering conditions remain limited^6,7^. Modern gene therapies present a compelling alternative, leveraging the inherent tropism of viruses to target specific cells and deliver therapeutic gene payloads for a range of spinal cord disorders (see ^8^ and ^9^ for review). Indeed, a number of viral gene therapies have progressed to human testing for spinal muscular atrophy^10–15^, giant axonal neuropathy^16^, neuronal ceroid lipofuscinoses type 7 disease^17^, and intractable cancer pain^18^. A variety of gene therapies have also been explored preclinically to target the cellular mechanisms of both spinal cord injury^19–41^ as well chronic/acute pain^42–53^. Among these, recombinant adeno-associated virus (AAV) vectors are particularly translatable owing to their replication deficiency and low risk of integration into the host genome^54^. While AAVs possess a comparatively robust viral safety profile (see ^55^ for review), these constructs can require an invasive intraspinal injection, which incurs risks of infection, blood vessel puncture, nerve damage, air embolism and disc penetration^56^ to bypass the molecularly restrictive blood-spinal cord barrier (BSCB) ^57,58^. To address these limitations, rationally designed AAV serotypes^59–63^ and administration of AAV to the cerebrospinal fluid^64,65^ have each been explored as substitutive means of transgene delivery to the spinal cord. In the absence of additional vector engineering, however, neither method provides the regional and cell-type specificity needed to preferentially express therapeutic transgenes in pathological spinal cord tissue. A developing technique, microbubble-aided focused ultrasound (FUS), has been shown by our group^66,67^ and others^68–70^ to be an effective alternative to surgical injection, allowing for noninvasive viral delivery to the central nervous system (CNS) via focal disruption of the neurovascular barriers regulating therapeutic access to these tissues.

Microbubble-aided FUS combines highly focused acoustic energy, which can penetrate bone and soft tissue, with intravenously administered microbubbles. At the focal target, the ultrasound induces microbubble oscillation, producing localized mechanical effects that transiently increase the permeability of neurovascular barriers, thereby permitting otherwise impermeant therapeutics to enter the neural parenchyma^71^. Although originally developed for molecule delivery across the blood-brain barrier (BBB) ^72^ microbubble-aided FUS has been demonstrated effective for spinal disruption of the rodent BSCB^73–78^ and has thereafter been translated to both rabbits^79^ and pigs^80^ (reviewed in ^81^). FUS BSCB disruption has been used to noninvasively deliver immunotherapeutics^82,83^, unencapsulated plasmid DNA^84,85^, and self-complementary AAVs^86^ to the rodent spinal cord/meninges, suggesting that this technique may be similarly suitable for viral gene therapies. Despite the considerable potential of these technical innovations, translation of FUS BSCB disruption to nonhuman primates - whose spinal cord physiology better approximates that of humans^87^ - remains an important step toward clinical deployment. Recently, we established safe parameters for FUS BBB disruption in the cortex of the common marmoset (*Callithrix jacchus*) ^88^. This development allowed for noninvasive delivery of AAV vectors encoding either fluorescent transgenes^67^ or excitatory chemogenetics^66^ within regions BBB disruption in marmoset cortex - providing a methodological foundation to extend FUS-mediated viral gene delivery to other regions of the marmoset CNS, including the spinal cord. Marmosets are well suited for evaluating FUS-based therapeutic strategies as they possess CNS architecture more similar to that of humans than other preclinical animal models^89–93^. Additionally, marmosets are amenable to transgenic modeling of diseases affecting both the brain^94,95^ and spinal cord^96–98^ allowing for a direct evaluation of treatment platforms targeting human pathologies. Of these, designer receptors exclusively activated by designer drugs (DREADDs) ^99^ are of particular interest for spinal cord pathologies due to their demonstrated clinical utility^100–104^, durability^105^, and ability to flexibly modulate cellular activity via systemic administration of an otherwise inert synthetic ligand (deschloroclozapine or DCZ) ^106,107^.

DREADDs have been widely implemented in the neurosciences since their inception nearly two-decades ago^99^ (see ^108^ for review). Following actuation via administration of a BBB/BSCB penetrant ligand, DREADDs can hyperpolarize (Gi) ^109^ or depolarize (Gq) ^110^ cells in which they are expressed. Though DREADDs are most well known in primates for their use in causal studies examining the neural bases of cognition and behavior (see ^111^ for a review of studies utilizing chemogenetics in nonhuman primates), this form of chemogenetics is also highly effective for the treatment of spinal cord injury^100,101^ and pain^102–104^ in preclinical animal models. DREADDs are also clinically advantageous as both their expression and function can be monitored *in vivo* using positron emission tomography (PET) ^66,106,112^. While these properties are encouraging for the burgeoning translation of DREADDs to the clinic, these constructs remain reliant on a direct viral injection for focal expression in the CNS^111^ (although see ^66^ and ^113^ for examples of FUS-based delivery of these constructs to the brain). Thus, establishing noninvasive - and spatially specific - DREADD delivery to spinal cord would provide both a translationally relevant therapeutic paradigm and a functional benchmark for viral gene delivery following FUS BSCB disruption - an approach that has, until now, remained largely unexplored in nonhuman primates.

Here, we develop the use of microbubble-aided FUS to noninvasively disrupt the BSCB for focal delivery of AAV vectors encoding either a fluorescent transgene or excitatory DREADDs within the marmoset spinal cord. By systematically testing a range of transducer central frequencies and ultrasonic pressures, we establish reliable parameters for robust molecular delivery within the marmoset cervical (C) and thoracic (T) spinal cord. We then apply these parameters for focal delivery of intravenously administered AAVs, either singly (AAV9-hSyn-hM3D(Gq)-mCherry) or as a pool (AAV9-CAG-GFP and AAV9-hSyn-hM3D(Gq)-mCherry), which transduced the spine within regions of BSCB disruption. To index the function of noninvasively transduced excitatory chemogenetics, we conducted FDG PET after systemic administration of the synthetic DREADD agonist, deschloroclozapine, ^106^ and found significantly elevated glucose metabolism in the region of FUS BSCB disruption as compared to untargeted spinal cord segments. Finally, we demonstrate the safety of FUS BSCB disruption in the marmoset though a combination of motor behavioral assay, gross anatomical assessment, and serial tape-transfer histopathological staining. Our results indicate that FUS BSCB disruption can be applied to nonhuman primates for safe, focal viral gene payload delivery. To facilitate broad application of this platform for noninvasive delivery of receptor-based gene therapeutics in marmosets, we generated an ultra-high-resolution (74 µm) multimodal MRI/CT spinal cord template for precise targeting and anatomical localization. We also provide open-access engineering drawings and CAD files for our M-FRAME system (Marmoset Fixation and Reorientation Apparatus for Multimodal Experiments), enabling precise and repeatable spinal targeting without surgical fixation (available at our Marmoset Neuroscientific Apparatus webpage - https://www.marmosetbrainconnectome.org/apparatus) ^114^. Together, these results establish FUS-mediated BSCB disruption as a safe and effective approach for noninvasive, focal gene delivery to the primate spinal cord.

## Results

### Noninvasive Disruption of the Marmoset BSCB

To establish FUS BSCB disruption within the marmoset spinal cord, we first sought to identify FUS parameters (central frequency and acoustic pressure) to reliably perturb the BSCB as indexed by extravasated volume of the MRI contrast agent gadolinium^88^. Previous reports have demonstrated that BBB disruption volume and intensity are related to ultrasonic central frequency and the peak negative pressure generated within tissue^88,115^. To probe for a similar relationship in the marmoset spinal cord, whose vertebral casing presents a vastly different bony geometry to the cranium, two marmosets received FUS using a 1.5 MHz or 550 kHz transducer each commanding either a higher (2.5 MPa, 1.5 MHz; 1.2 MPa, 550 kHz) or lower (1.8 MPa, 1.5 MHz; 0.75 MPa, 550 kHz) peak negative pressure (free-field). Immediately following application of FUS to the cervical (Marmoset V; one target at vertebral level C4) or thoracic (Marmoset S; 4 targets at vertebral levels T1, T5, T6, and T8) spinal cord, marmosets were transferred to an ultra-high field 9.4T MRI scanner and injected with 100 - 150 μL of a gadolinium-based contrast agent (GBCA) prior to MPRAGE image acquisition. Regions of BSCB disruption were identified and characterized via focal hyperintensities in MR images, which arise due to GBCA extravasation into targeted tissue. Figure 1 shows the target locations with accompanying central frequencies, acoustic pressures, and mechanical indices (MI) for each FUS target. Because of the small size of the spine, the possible resolution within our 86 mm volume coil was limited to 0.25 mm to acquire the scans within a reasonable time window that reduced the duration of anesthesia. To allow for better visualization of the anatomy of the spine, we also generated an ultra-high resolution MR marmoset spine template (see **Multimodal Marmoset Spine Template Generation**) from an *ex vivo* sample that was scanned over the course 72 hours. This template allowed for clear delineation of the internal structure of the cord at each target location to aid in localizing foci of BSCB disruption. GBCA images indicated successful BSCB disruption in all cervical (**Figure 1B**) and thoracic (**Figure 1C**) spinal targets (see **Figure 1A** for the rostral-caudal extent of BSCB disruption at each target on a template of the marmoset spine). Independent of peak negative pressure, we found that regions of BSCB disruption generated at 550 kHz (6.2 ± 2.4 mm^3^) were larger than those at 1.5 MHz (2.8 ± 0.1 mm^3^), as would be expected by the FWHM of the acoustic pressure distribution at the focus of each transducer (0.9 mm radial by 5.2 mm axial for 1.5 MHz and 2.4 mm radial by 16.8 mm axial radial for 550 kHz). The greatest volumes of BSCB disruption were achieved using a 550 kHz transducer commanding 1.2 MPa. These results suggest that a 550 kHz transducer commanding 1.2 MPa of pressure (free-field) maximizes the delivery of systemic agents within targeted regions of the spinal cord. More generally, our findings demonstrate that highly precise, noninvasive BSCB disruption is achievable in nonhuman primates at a variety of pressures and transducers. Our procedure implemented superiorly positioned, single element transducers, as we were able to rotate the marmoset torso (using our spine rotating apparatus, M-FRAME) to direct the ultrasound beam through spinal lamina rather than spinous processes. This approach dramatically simplifies the hardware necessary for FUS BSCB disruption in the marmoset.

**Figure 1.**
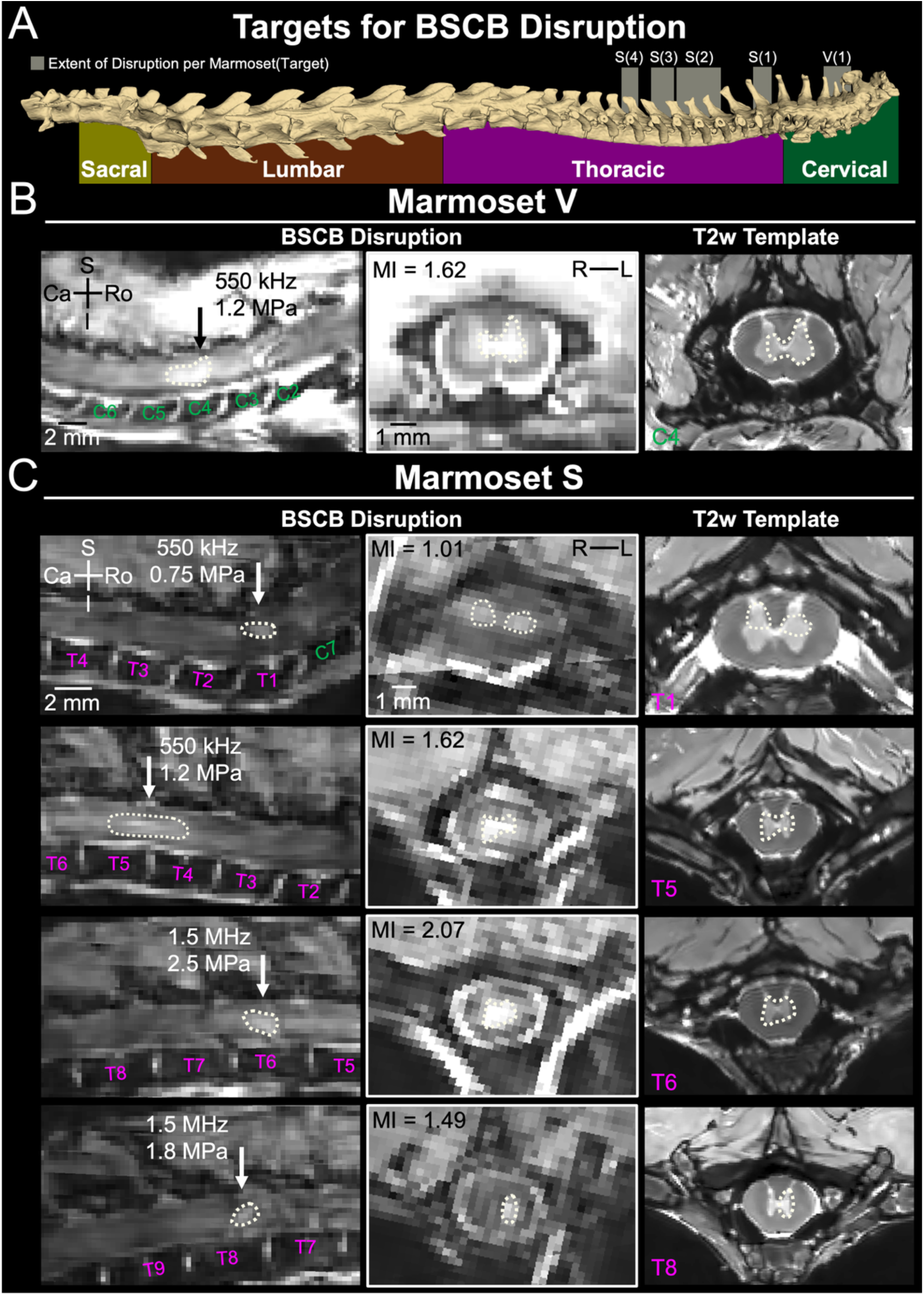
FUS BSCB disruption in the marmoset. **(A)** Template CT of the marmoset spinal cord. Semi-translucent boxes demarcate the approximate rostral-caudal extent of each region of BSCB disruption in Marmosets V and S. **(B, C)** Sagittal (left) and coronal (middle) MR images of BSCB disruption, as identified via focal hyperintensities arising from GBCA extravasation across the BSCB, in Marmosets V (B) and S (C). Arrows demarcate the position of corresponding coronal images along the rostral-caudal axis. FUS parameter sets (transducer central frequency and peak negative pressure (free-field)) for each treatment target are provided in sagittal images. Mechanical indices for each treatment target are provided in coronal images. MRI template images (right) are provided to visualize the internal anatomy of the marmoset spinal cord in relation to the extent of peak BSCB disruption (white dotted line) at the vertebral level of individual animal coronal images. Scale bars = 2 mm in sagittal images, 1 mm in coronal images. Abbreviations: S, superior; I, inferior; Ro, rostral; Ca, caudal; L, left; R, right; T, thoracic; C, cervical; MI, mechanical index

### Noninvasive AAV Delivery to the Marmoset Spinal Cord for Focal Transgene Expression

Having established FUS parameters to perturb the BSCB, we next combined FUS BSCB disruption with systemic AAV delivery. In these experiments, we applied FUS to the cervical (Marmoset H; 1 target at vertebral level C7/T1) and thoracic (Marmoset M; 1 target at vertebral level T2) spine using a 550 kHz transducer commanding a pressure of 1.1 MPa. Each target location and corresponding central frequency, acoustic pressure, MI and template image is shown in Figure 2. Immediately after verification of successful BSCB disruption via gadolinium-enhanced MPRAGE imaging (**Figure 2B, D**; see **Figure 2A** for the extent of BSCB disruption in both animals on a template marmoset spine), we administered AAVs either as a pool (AAV9-CAG-GFP and AAV9-hSyn-hM3D(Gq)-mCherry; Marmoset H) or singly (AAV9-hSyn-hM3D(Gq)-mCherry; Marmoset M) via an implanted venous catheter. Table 1 shows the construct(s), titer, volume and final dose administered to each animal. Both marmosets were harvested for verification of transgene expression by fluorescence microscopy four-weeks post-FUS application. In line with a previous report^86^, we found successful noninvasive transgene expression within all regions of BSCB disruption (in both Marmosets H and M), with minimal to no expression in other non-targeted portions of the spinal cord (**Figure 2C, E**). In Marmoset H, for example, very sparse glial GFP expression (regulated under the pan-cellular CAG promoter) was detected in an untargeted control section of the spinal cord (**Figure 2C, left**). This is likely due to our use of AAV9, which has some BSCB permeability in primates^116^. Within the region of BSCB disruption generated in Marmoset H, we found robust neuronal DREADD expression. GFP expression was also observed within both spinal neurons and glia (**Figure 2C**). Together, these results indicate that FUS BSCB disruption is an effective method for noninvasive, focal, cell type-specific delivery of both fluorescent and excitatory chemogenetic transgenes to the marmoset spinal cord.

**Figure 2.**
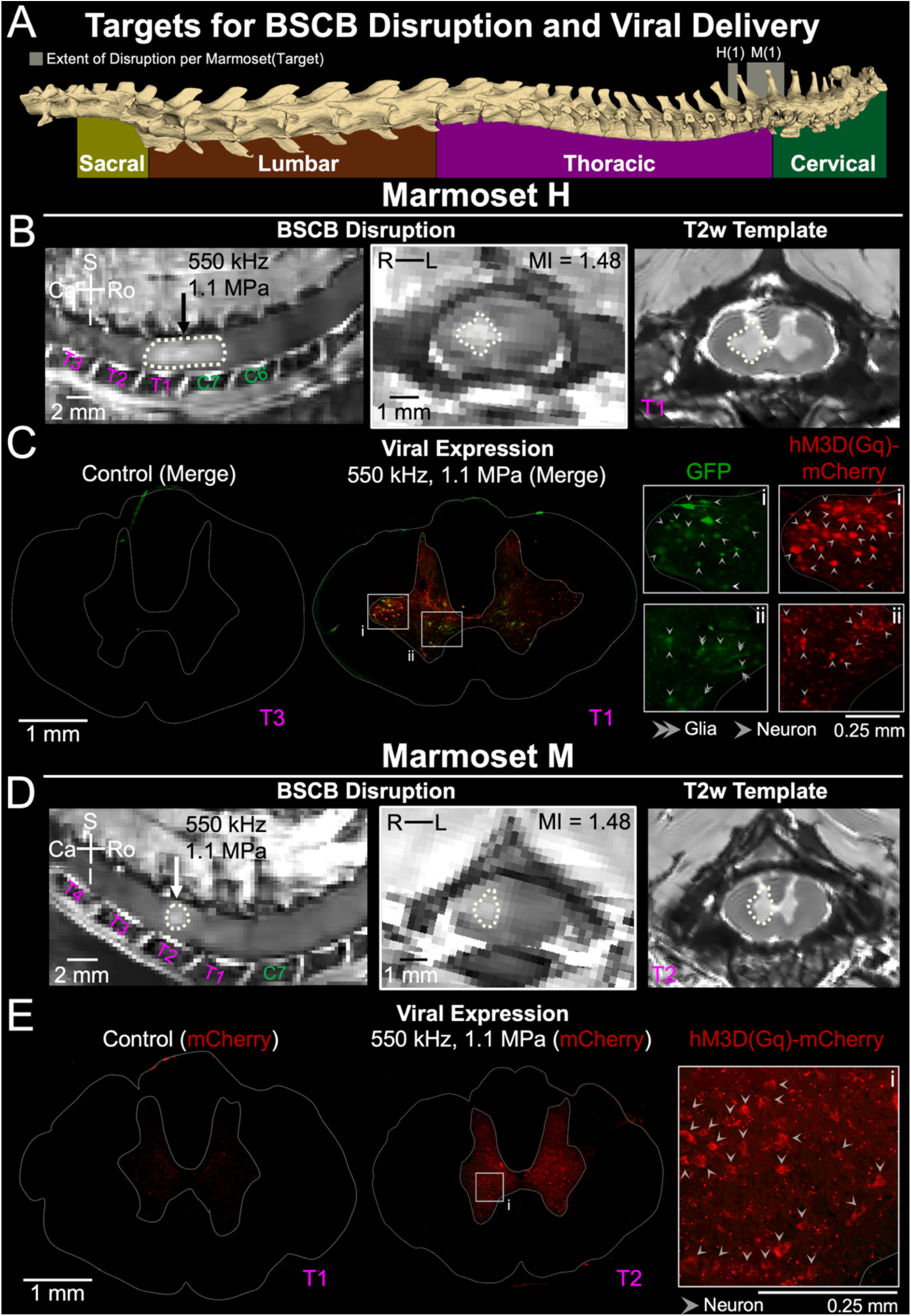
FUS BSCB disruption allows for noninvasive, focal viral delivery and transgene expression within the marmoset spinal cord. **(A)** Template CT of the marmoset spinal cord. Semi-translucent boxes demarcate the approximate rostral-caudal extent of BSCB disruption in Marmosets H and M. **(B)** Sagittal (left) and coronal (middle) MR images of BSCB disruption in Marmoset H. The arrow demarcates the position of the corresponding coronal image along the rostral-caudal axis. FUS parameters (transducer central frequency and peak negative pressure (free-field)) for the treatment target are provided in the sagittal image. The mechanical index of the treatment target is provided in the coronal image. An MRI template image (right) is provided to visualize the internal anatomy of the marmoset spinal cord in relation to the extent of BSCB disruption (white dotted line) at the vertebral level of the animal’s coronal image. **(C)** Fluorescence images of tissue sections sampled from a control region (left) and from within the area of BSCB disruption (middle) in Marmoset H, who received a systemic injection of a dual-AAV pool. Robust expression of both transgene constructs (hM3D(Gq)-mCherry and GFP) was observed within the region of BSCB disruption. Zoomed images of GFP and hM3D(Gq)-mCherry expression (right) correspond to regions demarcated in the tissue section sampled from the region of BSCB disruption (i, ii). Single arrows denote transgene expressing neurons while double arrows denote transgene expressing glia. **(D)** Same as (B) for Marmoset M. **(E)** Same as (C) for Marmoset M, who received a systemic injection of a single AAV encoding hM3D(Gq)-mCherry. Arrows denote transgene expressing neurons. Scale bars = 2 mm in sagittal MR images, 1 mm in coronal MR images, 1mm in tissue sections, 0.25 mm in zoomed images. Abbreviations: S, superior; I, inferior; Ro, rostral; Ca, caudal; L, left; R, right; T, thoracic; C, cervical; MI, mechanical index

**Table 1.**
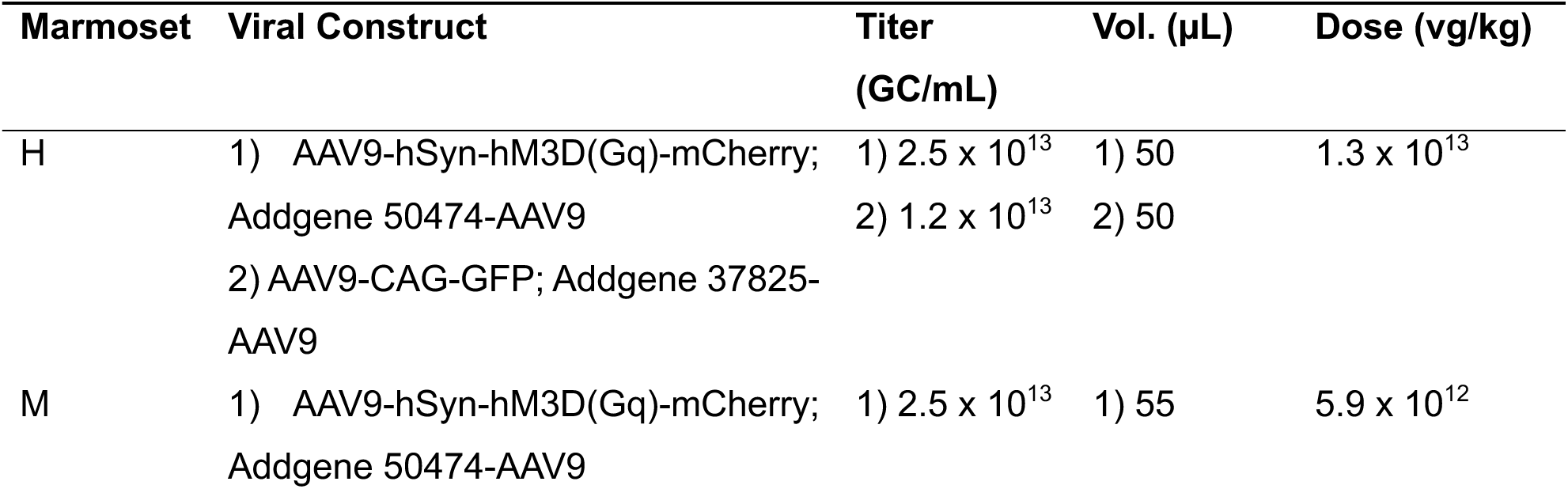
Details of post-FUS viral construct administration per marmoset.

### Regional Modulation of the Marmoset Spinal Cord via Noninvasively Transduced Excitatory Chemogenetics

To index the function of noninvasively transduced excitatory chemogenetics, we collected anesthetized FDG PET imaging data from Marmoset H following a period of 4 weeks to allow for adequate transgene expression^117^. PET data were collected immediately after intravenous administration of DCZ – a potent DREADD agonist. Following DREADD actuation, we observed significant increases in activation, measured by FDG uptake, within the region of BSCB disruption compared to other non-targeted segments of the spinal cord (**Figure 3A, B**) as indicated by a Wilcoxon signed rank test corrected for multiple comparisons (T1/C7 (region of BSCB disruption) vs C6/C5, adjusted *p* < 0.001, T1/C7 vs C4/C3, adjusted *p* < 0.001; T1/C7 vs T3/T2, adjusted *p* < 0.001). Visual examination of FDG PET data suggested the presence of a rostrally spreading depolarization from the region of BSCB disruption. Statistical analysis confirmed this observation, indicating that, external to the region of BSCB disruption, FDG uptake was greatest in the cervical spine directly adjacent to the region of BSCB disruption and that uptake decreased rostrally from the site of transgene expression (Wilcoxon signed rank test corrected for multiple comparisons, C6/C5 vs C4/C3, adjusted *p* < 0.001; C6/C5 vs T3/T2, adjusted *p* < 0.001; C4/C3 vs T3/T2, adjusted *p* < 0.001). These data indicate that excitatory chemogenetics expressed within the region of BSCB disruption can be functionally modulated.

**Figure 3.**
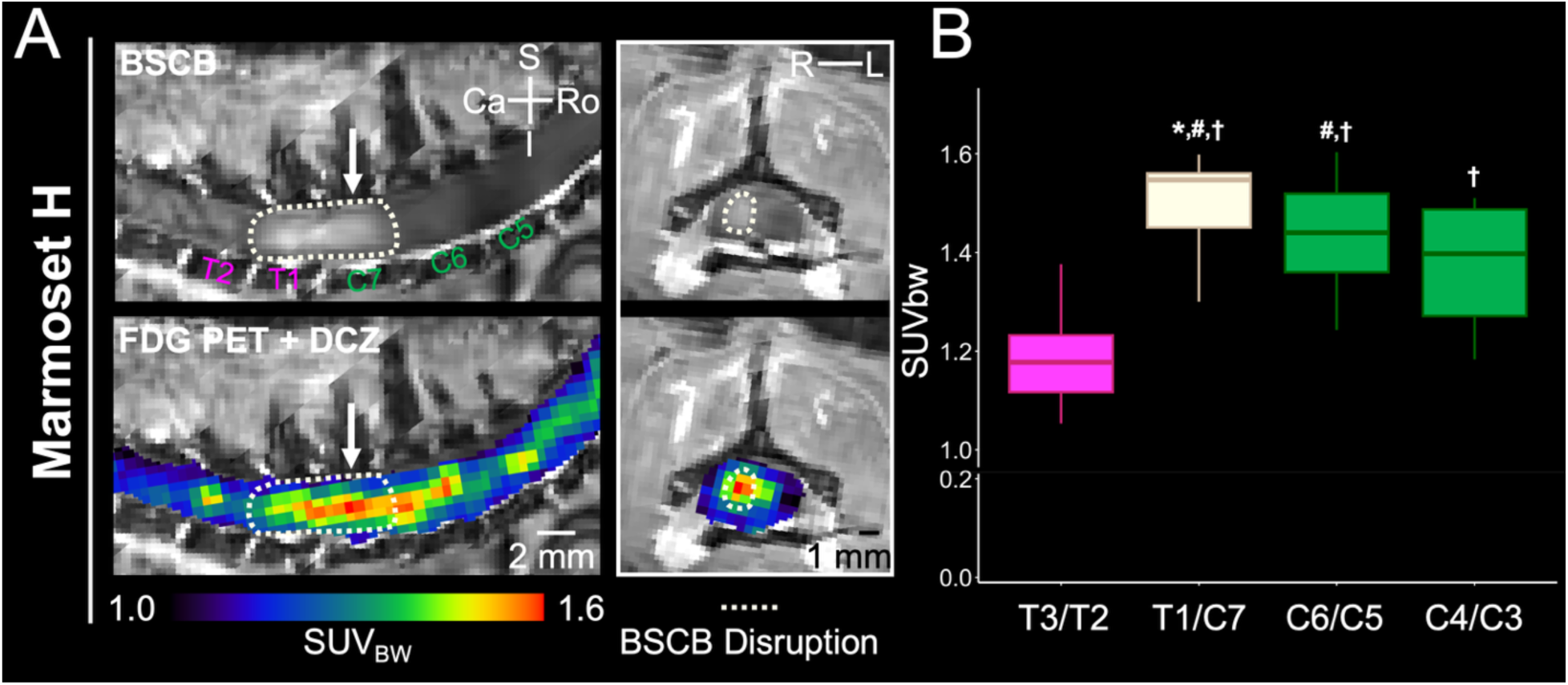
DREADD actuation of noninvasively transduced chemogenetics increases FDG uptake within the region of BSCB disruption and in rostral spinal segments. **(A)** Row 1 shows the extent of BSCB disruption (white dotted line) in the sagittal (top left) and coronal (top right) planes for Marmoset H. Row 2 shows weight normalized FDG uptake following actuation of noninvasively transduced chemogenetics in the sagittal (bottom left) and coronal (bottom right) planes. **(B)** Box plot of weight-normalized FDG uptake (median and interquartile range) in the region of BSCB disruption (white, T1/C7) as well as in caudal (T3/T2, pink) and rostral (C6/C5), C4/C3, green) spinal segments. A Wilcoxon signed rank test indicated that FDG uptake following chemogenetic actuation was greatest in the region of BSCB disruption (T1/C7 (region of BSCB disruption) vs C6/C5, adjusted *p* < 0.001, T1/C7 vs C4/C3, adjusted *p* < 0.001; T1/C7 vs T3/T2, adjusted *p* < 0.001). Actuation of DREADDs expressed within the region of BSCB disruption also increased FDG uptake rostrally (C6/C5 vs C4/C3, adjusted *p* < 0.001; C6/C5 vs T3/T2, adjusted *p* < 0.001; C4/C3 vs T3/T2, adjusted *p* < 0.001). Scale bars = 2 mm for sagittal images, 1 mm for coronal images. † greater than T2/T3; * greater than C6/C5; # greater than C4/C3 Abbreviations: S, superior; I, inferior; Ro, rostral; Ca, caudal; L, left; R, right; T, thoracic; C, cervical

### Multimodal Safety Assessment of FUS BSCB Disruption

It has been shown that FUS application to the rodent spinal cord can deleteriously affect motor function^73,78,86^. To probe for motor deficits following FUS application to the marmoset spinal cord, we employed an assessment of upper limb function commonly used in marmoset stroke models known as the Valley Task^118^. Valley Task testing was conducted in Marmoset H (during the period viral expression post-FUS application) as the region of BSCB disruption in this animal covered the caudal portion of the brachial spinal cord innervating the arm (**Figure 2B**) ^119^. In the Valley Task, marmosets use one arm to collect food rewards positioned at ascending levels of elevation (difficulty) via a centrally located opening in the testing apparatus. Rewards are placed at either the right or left extreme of the apparatus which allows for an assessment of motor performance in each arm separately over the course of a testing session (**Figure 4A**). Two metrics were calculated for each limb: a total accuracy score and the latency to collect all food rewards. Linear mixed-effects models were fit to behavioral data to examine fixed effects of time (days post-FUS), arm (left vs right) and their interaction on Valley Task score and latency. For each model, the testing session was held as a random effect. We observed no main effects of time post-FUS or hand for Valley Task score (time, *F*(1,11) = 0.82, *p* = 0.38; arm, *F*(1,37) = 0.62, *p* = 0.44). The interaction between time post-FUS and arm was also not significant for Valley Task score (F(1,37) = 0.05, *p* = 0.83). No significant effect of arm was found for latency (*F*(1,37) = 0.63, *p* = 0.43). We did, however, identify a significant main effect of time post-FUS for latency (*F*(1,11) = 28.99, *p* < 0.001) suggesting that Marmoset H completed the test faster as time progressed (**Supplemental Figure 1**) but the interaction between time post-FUS and arm was not significant (*F*(1,37) = 0.15, *p* = 0.70). Indeed, differences between the left and right hand for both Valley Task score and latency remained minimal over the course of behavioral testing (**Figure 4B**). These results suggest that FUS application to the marmoset spinal cord elicits no long-lasting motor deficits, which would be relegated to the right arm/hand based on the unilateral position of BSCB disruption. Although we did detect DREADD expression within this animal, we did not collect behavioral data after DREADD actuation due to the potential for pan-neuronal DREADD expression (including cells mediating paint) conferred by the promoter used in this study.

**Figure 4.**
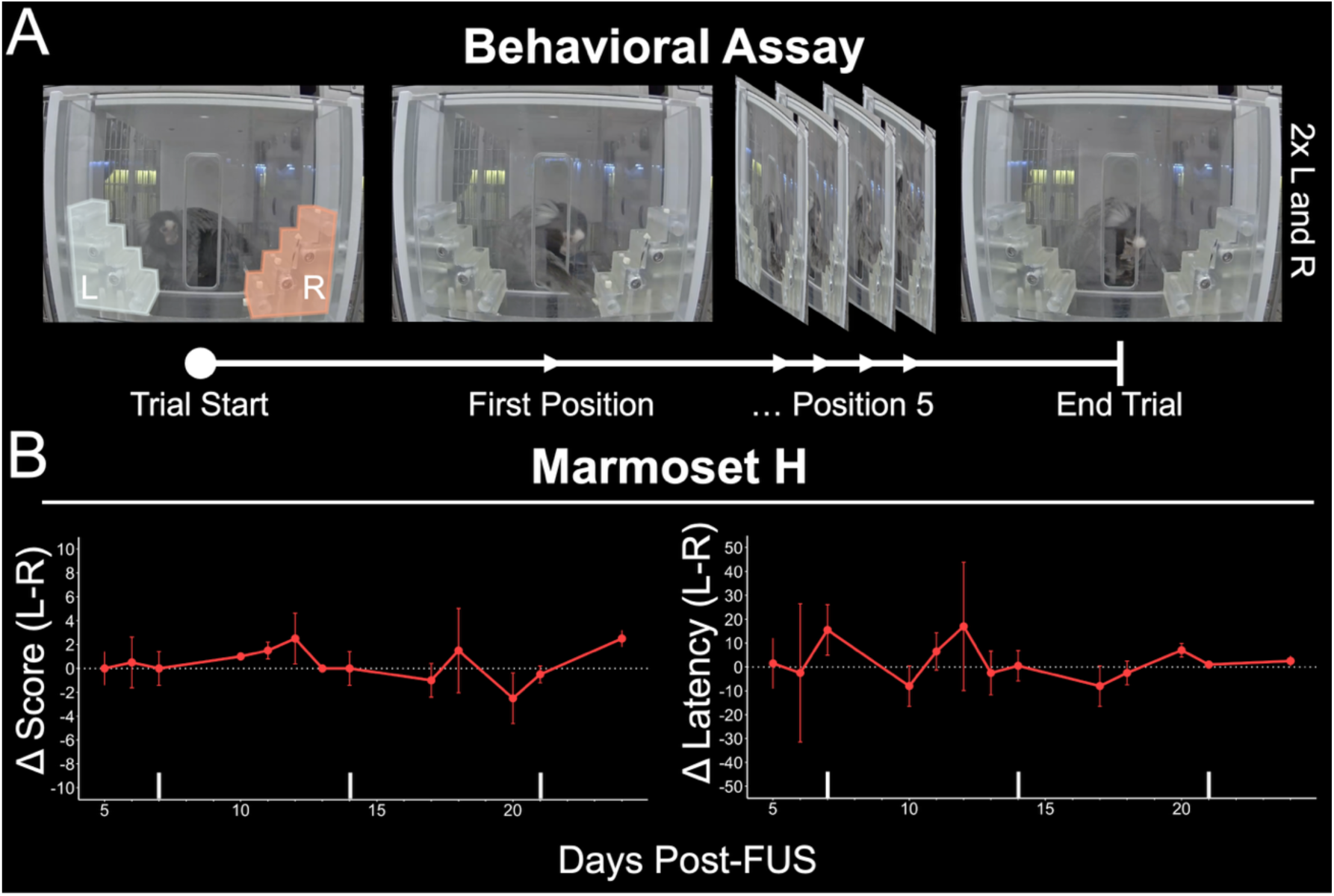
FUS BSCB disruption does not decrease upper limb motor performance in the marmoset. **(A)** Overview of the Valley Task^118^ as an assessment of post-FUS upper limb motor function. Marmosets were required to collect food rewards placed at ascending levels of the lateral extremes of the testing apparatus^114^ using either the left or the right arm. Two trials for the left and right arm were collected during each session. Behavioral data was collected from Marmoset H, as the region of FUS BSCB disruption in this animal included the brachial portion of the spinal cord which innervates the upper limb^119^. Two metrics, stratified by arm, were calculated from behavioral data: Valley Task score and latency. **(B)** Difference in Valley Task score and latency (mean ± SD) between the left and right arm over successive behavioral sessions collected between five and 24-days post-FUS. The region of BSCB disruption was restrained to the right portion of the spinal cord, which corresponds to the right arm. Valley Task score and latency were comparable for each arm across the entire duration of testing.

Although we did not detect motor behavioral deficits following FUS BSCB disruption, it is possible that this technique generates damage only detectable via dedicated *ex vivo* analyses. To probe for FUS-mediated tissue damage, we extracted all spinal cords at the conclusion of experiments for both a gross anatomical assessment (Marmosets V, H and M) as well as histopathological staining (Marmosets V and S). Extracted spinal cords displayed no external signs of FUS-induced hemorrhage (**Figure 5A**; see refs. ^80^ and ^88^ for examples of such tissue damage in the pig spinal cord and marmoset brain, respectively). In complement, we found no signs of microscopic damage in histochemically stained, serial tissue sections from Marmosets V and S (**Figure 5C, D**). Specifically, examination of tissue sections from within regions of BSCB disruption revealed no evidence of extra-vessel erythrocyte accumulation (H&E staining), overt tissue tearing, abnormal cell morphology, and unhealthy cell nuclei (Nissl staining). These data, in combination with those from our post-FUS motor behavioral assessment, provide compelling evidence that FUS BSCB disruption within the nonhuman primate spinal cord is both behaviorally and mechanically safe.

**Figure 5.**
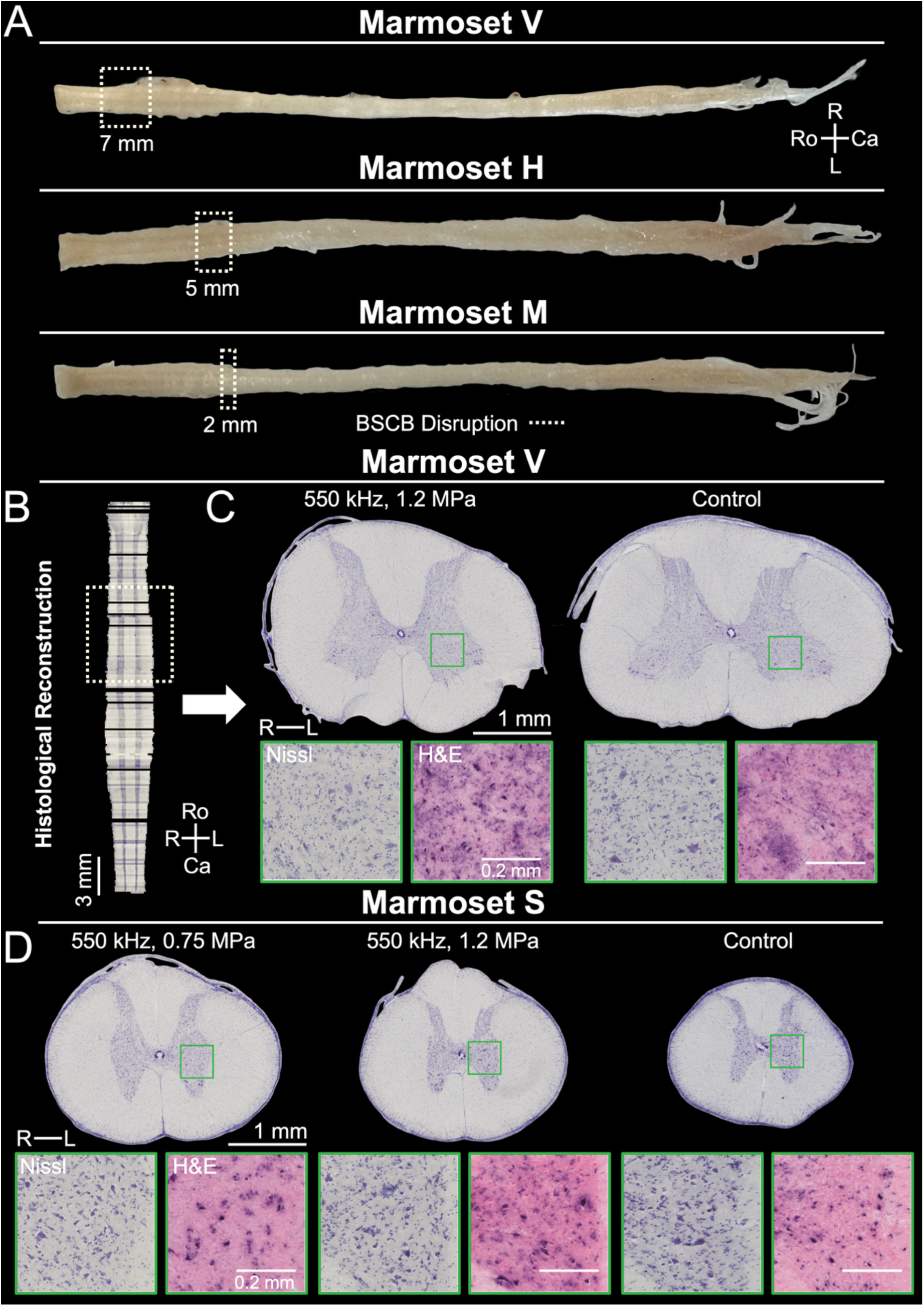
FUS BSCB disruption does not damage the marmoset spinal cord. **(A)** Extracted spinal cords from marmosets V, H and M. **(B)** Example of volumetric reconstruction using Nissl stained sections from Marmoset V, which aids in identifying tissue sections within the region of BSCB disruption. For both the extracted spinal cords and the volumetric reconstruction, white boxes demarcate the extent of BSCB disruption. **(C)** Representative Nissl stained sections from Marmoset V sampled from within the region of BSCB disruption (left) and an external control region (right). **(D)** Representative Nissl stained sections from Marmoset S sampled from within regions of BSCB disruption (left, middle) and an external control region (right). Images below each tissue section in (C) and (D) depict zoomed in views of Nissl and H&E staining within the corresponding green ROI. Labels above each section provide the FUS parameters used to generate the region of BSCB disruption from which non-control sections are taken. Scale bars = 1 mm for Nissl sections, 0.2 mm for zoomed images. Abbreviations: Ro, rostral; Ca, caudal; L, left; R, right

## Discussion

Gene therapies hold great promise to treat the cellular mechanisms directly contributing to spinal pathology (see ^8^ and ^9^ for review). Despite marked progress in using viral-based gene delivery systems as therapies for in preclinical models of disease^19–53^, several challenges remain in translating these findings to human patients. Principal among these is the requirement for highly invasive procedures (laminectomy, intraspinal injection, etc.) to promote focal transgene expression in pathological tissue. Here, our goal was to establish FUS BSCB disruption for noninvasive, focal delivery of neurotropic viruses encoding excitatory chemogenetics to the nonhuman primate spinal cord. We first established parameters for FUS BSCB disruption in the marmoset to allow penetrance of systemically administered molecules within discrete segments of the spinal cord. We then showed that FUS BSCB disruption allows for focal transduction of excitatory chemogenetic and fluorescent transgenes within neuronal and glial cells in the targeted region of the spinal cord. To verify that noninvasively transduced chemogenetics remained functional in the marmoset spinal cord, we conducted FDG PET imaging immediately following chemogenetic actuation and found increased metabolic demand within the region of BSCB disruption. Confirming the safety of FUS-mediated BSCB disruption in the brachial region of the spinal cord, no deficits were observed in longitudinal post-FUS motor behavior assessed using the Valley Task. These findings were corroborated by gross *ex vivo* examination of the spinal cord and serial tape-transfer histology, which enabled staining of every section throughout the cervical and thoracic spinal cord and revealed no evidence of tissue injury. These findings outline a promising platform for noninvasive, safe viral gene therapeutic delivery and testing within the nonhuman primate spinal cord. To assist in continued development of FUS-based gene therapeutic delivery in marmosets, we provide public access to our high-resolution, multimodal template of the marmoset spinal cord as well as our hardware platform for marmoset spine FUS and anatomical imaging, M-FRAME, via our Marmoset Neuroscientific Apparatus webpage (https://www.marmosetbrainconnectome.org/apparatus/).

Most extant studies evaluating FUS BSCB disruption have been conducted in rodents, with comparatively limited work in other mammalian preclinical models (see ^81^ for review). We therefore sought to establish parameters for noninvasive FUS BSCB disruption in marmosets with the express intent of maximizing focal AAV delivery to the spinal cord in nonhuman primates. Two of the five marmosets included in this study received FUS to the cervical (Marmoset V, **Figure 1B**) and thoracic (Marmoset S, **Figure 1C**) spinal cord for the purpose of testing across a range of ultrasonic pressures and frequencies. As expected, we found that the size of disruption increased when employing either a lower frequency ultrasound transducer or when commanding a high peak negative pressure within targeted tissue. The largest regions of BSCB disruption, which were generated by the 550 kHz transducer commanding 1.2 MPa, remained spatially constrained within a maximum of ∼4 mm (one vertebral level) from the targeted site. Our findings identified 550 kHz at a high (1.2 MPa) peak negative pressure (free-field) as a favorable parameter combination for noninvasive viral delivery to the spinal cord, producing extensive but regionally limited BSCB disruption at a comparatively lower - and FDA approved (< 1.9 for diagnostic imaging) ^120^ - MI of 1.62 (non-derated). Although use of a lower frequency transducer permitted command of reduced peak negative pressures for robust BSCB disruption, both the higher and lower pressure levels presented here remain elevated as compared to those commonly employed for rodents of comparable size to marmosets (e.g., rats) ^74,75,77,82,83^. For example, O’Reilly et al. (2018) ^82^ utilized a spherically focused 551.5 kHz transducer commanding a mean peak pressure of 0.43 MPa to disrupt the rat BSCB barrier – a value nearly half that of the lowest pressure generated by our 550 kHz transducer. It therefore remains possible that BSCB disruption in the marmoset is achievable at lower pressures than those presented here. Comparisons between parameter sets necessary for BSCB disruption in marmosets and other non-primate species are further complicated by the varying physical properties of ultrasound transducers employed across labs and tissue through which the beam must pass to generate pressure within the spinal cord. As such, the FUS parameters presented here represent conditions sufficient to support delivery of systemically injected agents within the marmoset spinal cord, not minimum thresholds for FUS BSCB disruption.

Previous work in rodents suggests that FUS BSCB disruption may be suitable for delivery of non-receptor-based viral gene therapeutics^86^. Engineered receptor-based systems provide an added clinical benefit in that their actuation is dependent upon administration of nonendogenous signaling molecules which allows for noninvasive, reversible therapeutic modulation. To examine whether such systems could be delivered noninvasively to the marmoset spine, we applied our optimized FUS parameters to promote regional penetrance of AAV vectors encoding either excitatory DREADDs or fluorescent transgenes within the spinal cord. Two marmosets received FUS targeting either the cervical (Marmoset H, **Figure 2B**) or thoracic (Marmoset M, **Figure 2D**) spinal cord using a 550 kHz transducer commanding 1.1 MPa. Following MRI-based verification of BSCB disruption, we systemically administered a bolus of AAV either singly (AAV9-hSyn-hM3D(Gq)-mCherry, Marmoset M) or as a pool (AAV9-hSyn-hM3D(Gq)-mCherry and AAV9-CAG-GFP, Marmoset H) via an implanted venous catheter. Both single and dual-AAV administration post-FUS resulted in robust transgene expression within the region of BSCB disruption (**Figure 2C, E**). We also observed limited - and much less intense - transgene expression elsewhere in the spinal cord, likely owing the use of AAV9 which has been shown to transduce spinal motor neurons in macaques following systemic administration^116^. Additionally, Weber-Adrian et al. (2015) ^86^ have shown that systemic administration of a high AAV dose (7 x 10^12^ vg/kg) following disruption of the rat BSCB results in transgene expression contralateral to targeted tissue. Like in the brain, other serotypes will likely allow for even more exclusive passage across the BSCB. Within regions of BSCB disruption, we observed robust neuronal DREADD expression across both animals. In Marmoset H, who received a systemic injection of a dual-AAV pool, we also observed GFP expression in both spinal neurons and glia. We note that although the volume of each construct composing the pool was the same, the titer of the DREADD encoding construct was greater than that encoding GFP, likely accounting for the (slight) differences in transduction efficacy^121^. Together, these results demonstrate proof-of-principle for noninvasive, focal delivery of potential receptor and non-receptor based viral gene therapeutics within the primate spinal cord either singly or combined in a dual-AAV pool.

To index the function of noninvasively transduced DREADDs in the marmoset spine, we conducted FDG PET imaging in Marmoset H following intravenous injection of the synthetic DREADD agonist, DCZ. We found that chemogenetic actuation significantly increased metabolic demand in the region of BSCB disruption as compared to other untargeted spinal segments (**Figure 3**). We also observed a gradient of increased activation extending rostrally from the site of chemogenetic expression (**Figure 3**). These results are in line with our previous report that actuation of noninvasively expressed chemogenetics within regions of BBB disruption modulates activation both locally (∼4 mm from the site of transgene expression) and across entire frontocortical circuits^66^. The presence of spreading or, perhaps, circuit wide depolarization from the region of spinal expression following chemogenetic actuation is a key consideration for therapeutic translation. In mice, for example, actuation of excitatory DREADDs targeted to lumbar projecting propriospinal neurons of the thoracic spinal cord significantly improves hindlimb locomotor function following complete paralysis – a clearly beneficial therapeutic effect. ^100^ Conversely, however, chemogenetic modulation of rodent sensory circuits can reduce sensation of innocuous stimuli – a potentially detrimental side effect^53^. Although clinical sequalae of spreading modulation following chemogenetic actuation remain to be tested, these results demonstrate the utility of FUS BSCB disruption for noninvasive delivery of highly translational neural actuators to the nonhuman primate spinal cord.

Procedure related motor deficits^73,78,86^ and tissue damage^74,75,79,80,86^ have been reported following FUS BSCB disruption. One challenge in developing FUS for safe BSCB disruption arises from the irregular bony geometry of the vertebral column. Spinous processes distort and absorb the ultrasound beam while the vertebral body reflects sound, generating an irregular pressure distribution (standing waves) and regions of substantially increased ultrasonic pressure (hotspots) ^79^. To eliminate confounds arising from the spinous processes, a number of labs have adopted a pre-FUS laminectomy^77,79,84^. Others have employed FUS apparatus that incorporates two transducers positioned on either side of the spinous processes which also enables the use of short-burst, phase-keying to attenuate standing wave formation^74,75,80^. For our study, we elected to utilize single element transducers and devised an apparatus enabling rotation of the marmoset torso eliminating the need for an invasive laminectomy to avoid beam disruption from the spinous processes. To evaluate the neurological safety of this alternative method of FUS BSCB disruption, we conducted a motor behavioral assay of the upper limb in Marmoset H and found no difference in limb performance following BSCB disruption within the right brachial spinal cord (**Figure 4**). Indeed, the latency to complete the task decreased over time but not in a manner that was arm dependent (**Supplemental Figure 1**) as would be expected following unilateral damage to the spinal cord. We note that, although this animal did express excitatory DREADDs within the spinal cord (**Figure 2C**), we chose not to collect behavioral data following chemogenetic actuation to avoid potentially untoward effects from broad neuronal transduction (including cells that could cause pain). To further assess the safety of FUS BSCB disruption, extracted spinal cords from Marmosets V, H, and M were examined for FUS induced tissue damage. We observed no tissue tearing or hemorrhage within the region of BSCB disruption in extracted spinal cords (**Figure 5A**, see refs. ^80^ and ^88^ for examples of macroscopic FUS-induced tissue damage in the spinal cord and brain, respectively). To probe for microscopic damage, we conducted serial tape-transfer histochemical staining on tissue from Marmosets V and S which showed no evidence of FUS-induced microbleeds or abnormal cell/tissue health within regions of BSCB disruption (**Figure 5B, C**). These data suggest that neurologically and mechanically safe BSCB disruption is possible in marmosets using a single element transducer of either high (1.5 MHz) or low (550 kHz) frequency and simple rotation of the torso. Although we cannot rule out the possibility that our preparation could result in standing wave formation within the marmoset spinal cord, our data suggest that potential regions of elevated ultrasonic pressure were insufficient to generate tissue damage. Even so, incorporation of an additional transducer to implement a short-burst, phase-keying pulse sequence presents a compelling option for further technical development in marmosets but is not strictly necessary for safe, precise, laminectomy-free FUS BSCB disruption and viral delivery in this nonhuman primate species.

An emerging concern in the field of viral gene therapy is the large viral dose required for systemically injected vectors to penetrate the neurovascular barriers protecting the CNS. At sufficiently large doses, systemic AAV injection may initiate deleterious inflammation as well as liver and dorsal root ganglion toxicity^122^. Although FUS BSCB disruption can effectively lower the required viral dose for robust transduction within targeted regions in rodents^86^, the doses presented here remain elevated compared to those that would typically be employed for an intraparenchymal injection. AAV vector engineering presents an intriguing option to reduce the viral dose required for robust transgene expression within regions of BSCB disruption and attenuate off target peripheral organ accumulation. A recently developed AAV9 variant known as AAV.FUS.3, for example, displays enhanced tropism for cells within regions of BBB disruption and is detargeted from the murine liver (as compared to AAV9) ^123^. Dedicated testing of these variants in nonhuman primates will be critical in attempts to reduce viral doses, however, as the optimized tropism of engineered vectors can be species specific^124^. Nevertheless, we detected no clinically apparent adverse health events in the animals that underwent the FUS BSCB disruption and viral delivery experiments reported here. Moreover, the largest viral dose we employed was much lower than that used for diffuse BSCB penetrance in other nonhuman primates^116^ and approximately one order of magnitude smaller than those employed in human clinical testing^10–16^.

In summary, we demonstrate that safe, noninvasive, focal BSCB disruption is achievable in marmoset nonhuman primates and that this technique enables targeted AAV delivery across the BSCB for chemogenetic and fluorescent transgene expression. Using *in vivo* PET imaging, we further establish that noninvasively transduced excitatory chemogenetics are functional in the marmoset spine. We also provide both our M-FRAME hardware platform and ultra-high resolution multimodal marmoset spine template publicly to support continued development of FUS in the primate spine. Together, these resources and findings compose a platform for systematic evaluation of targeted viral gene therapies that is simpler, safer, and less expensive than conventional surgical methods for AAV delivery to the spinal cord. With proof-of-principle for noninvasive DREADD expression demonstrated here, this technique could be leveraged for a variety of receptor-based constructs, thereby allowing controlled, therapeutic stimulation of spinal circuits across a variety of disease states at scales necessary for wide clinical deployment.

## Methods

### Animals

Five adult marmosets (*Callithrix jacchus*; *n* = 3 female) contributed data to this study (**Supplementary Table 1**). Prior to survival procedures (FUS, MRI and PET/CT), marmosets received intramuscular injections of meloxicam (0.2 mg/kg; 07-893-5916, Patterson Vet Supply, Inc., Loveland, CO, USA) and glycopyrrolate (0.005 mg/kg; 1226607, McKesson Medical-Surgical, Richmond, VA, USA). All animals were anesthetized (induced and maintained) with 2% isoflurane delivered via mask for the FUS, MRI, PET and CT procedures. During the procedures, heart rate, blood oxygenation, respiration, and rectal temperature were monitored. For FUS experiments, the back was shaved with clippers, then any remaining hair was removed with depilatory cream. A 26-gauge catheter was placed in the lateral tail or saphenous vein for microbubble and contrast agent delivery. Body temperature was maintained with heated water blankets. Experimental procedures were approved by the University of Pittsburgh Institutional Animal Care and Use Committee.

## Apparatus Generation

### In Vivo MRI, PET/CT Imaging Apparatus

To optimize spine imaging in both PET/CT and MRI applications (as described below), we designed an experimental platform termed M-FRAME (Marmoset Fixation and Reorientation Apparatus for Multimodal Experiments) that allows for imaging in the prone position, while meeting spatial constraints across small imaging bores (as small as 86 mm inner diameter) (**Figure 6**). The primary purpose of this system is to image the spine with minimal curvature across the rostral-caudal and medial-lateral axes. Straightening the spine in this way reduces MRI signal inhomogeneity while also simplifying future spatial comparisons and post-imaging alignment across modalities (MRI, CT, PET, the FUS stereotactic system, and fixed tissue). With a pitch-rotated anesthesia mask, the marmoset can be mounted with ear bars in the standard sphinx position, then the head rotated downwards to minimize curvature of the cervical spine - a technique we have successfully performed in Cebus and macaque monkeys over hours-long MRI sessions^125^. All computer-aided design (CAD) models were designed in SolidWorks (version 2022, Dassault Systèmes SolidWorks Corporation, Waltham, MA, USA), with Initial Graphics Exchange Specification (IGES), Standard for the Exchange of Product Model Data (STEP), and stereolithography (STL) files made publicly available for download through our Marmoset Neuroscientific Apparatus webpage (https://www.marmosetbrainconnectome.org/apparatus/). These models were designed to minimize thickness for PET/CT signal attenuation, while also being MRI-compatible at ultra-high field, producing minimal susceptibility artifact^125^. With these considerations in mind, two 3D printed materials were used: acrylonitrile butadiene styrene (ABS) or nylon 12 powder (Nylon 12 Powder, FUSE 1+ 30W, Formlabs, Somerville, Massachusetts, USA).

**Figure 6.**
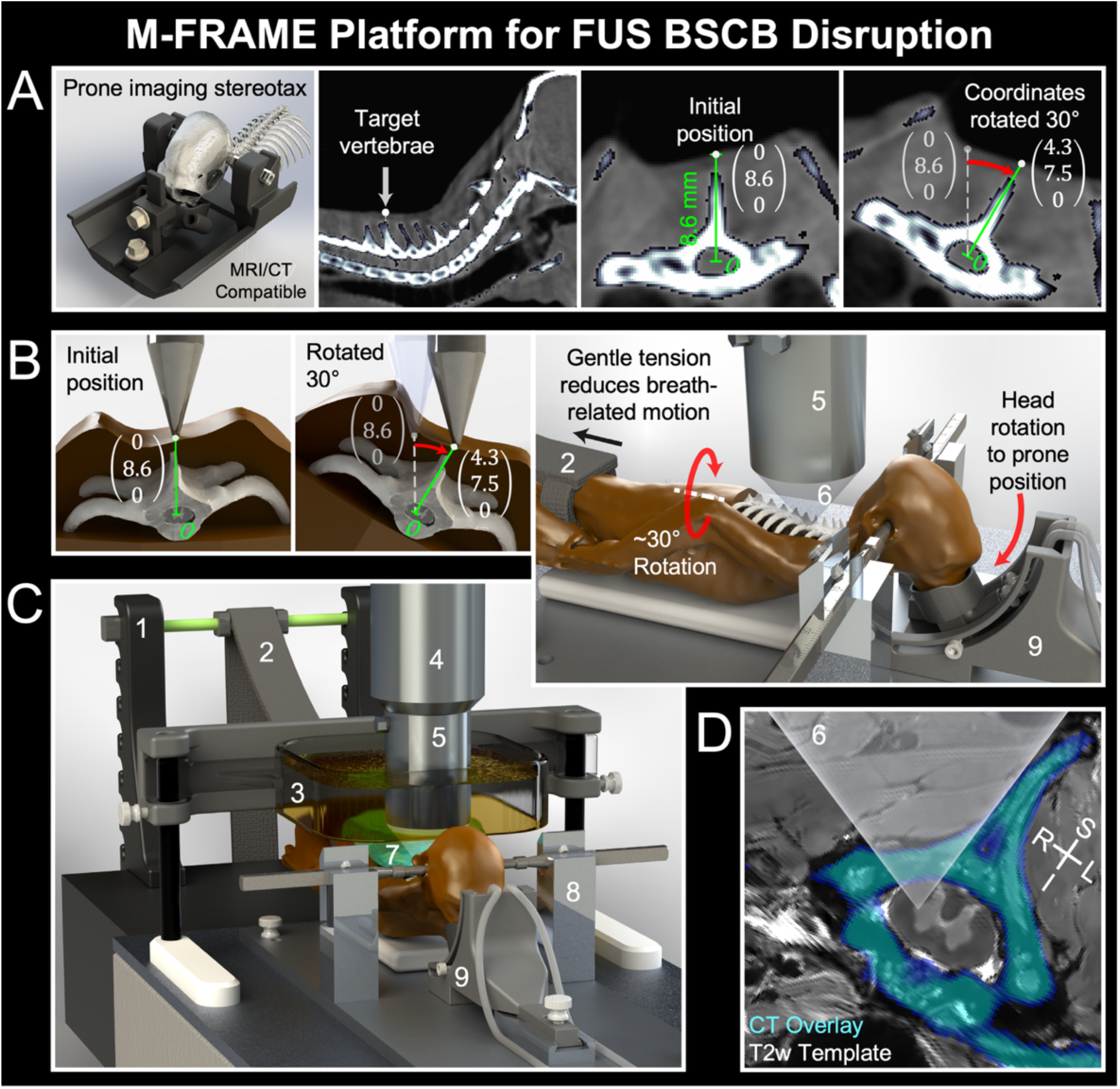
M-FRAME hardware platform for FUS BSCB disruption and structural imaging. **(A)** MRI/CT compatible prone stereotax (left) for structural imaging of the marmoset spinal cord. This stereotax was employed for pre-FUS CT scanning, as it allows for image acquisition while the marmoset is in a position similar to that for FUS BSCB disruption (second from left). Coordinates of FUS targets were identified in the pre-procedural CT scan (second from right) and rotated ∼30° counterclockwise via a matrix calculation (right, **Equation 1**) to maintain targeting after rotation of the marmoset torso via the FUS component of M-FRAME. **(B)** Targeting strategy for FUS BSCB disruption (left, middle). For FUS BSCB disruption, the M-FRAME platform incorporates a rotatable anesthesia mask to pitch the head downward and allow transducer access to the cervical spine. This system also utilizes two hook-and-loop strap systems to apply gentle caudal pressure to the hips, thereby straightening the spine and eliminating motion from respiration, and to rotate the torso ∼30°, which directs the ultrasound beam through the spinal lamina rather than the spinous process (right). **(C)** Schematic of the M-FRAME platform assembled for BSCB disruption. **(D)** Visual representation of our rotational targeting strategy in MRI and CT template images of the marmoset spine. Rotation of the marmoset torso directs the ultrasound beam (semi-translucent cone) through the lamina of the vertebra (cyan overlay). The number labels in (B), (C) and (D) of this figure correspond to the following: 1) Hook style mounts for hip suspension; 2) Fabric strap for hip suspension; 3) Degassed water bath; 4) Transducer attachment; 5) Ultrasound Transducer; 6) Ultrasound beam; 7) Ultrasound gel for acoustic coupling to the back; 8) Stereotax; 9) Rotatable anesthesia mask. Abbreviations: S, superior; I, inferior; L, left; R, right

### FUS Apparatus for BSCB Disruption

Due to the complex bony geometry of the spine and the small size of the spinal cord in small animal models, precise FUS delivery to the spinal cord has been achieved with an invasive stereotactic apparatus^77^ or though imaging guidance^86^ – both of which present an issue for the translatability (invasive, contacting bone or cost with MRI) of the FUS BSCB disruption technique. As such, we developed an apparatus that takes a simpler, noninvasive approach utilizing a single, spherically focused FUS transducer and a pre-procedural CT scan for treatment planning (**Figure 6B, C**). There were several considerations in the design; first, with the marmoset in the sphinx position, the head occludes the cervical spine from above, complicating FUS delivery to the cervical enlargement – this issue was resolved by rotating the head into the prone position. Second, with the animal in the prone position, respiration changes the position of the spine due to torso contact with the stereotax base – we resolved this through gentle hip suspension by way of an adjustable hip strap apparatus consisting of 3D-printed hook-style mounts (ABS), a quarter-inch fiberglass rod fulcrum, and a fabric strap with hook-and-loop fasteners. This served to not only suspend the hips, but to also straighten the spine. Third, we implemented a simple mechanical targeting system to leverage the unique morphology of the spinous processes. With these processes visible through the skin and identifiable through their distinct shape – for example with segment C7 having a taller process than C6 – we could identify and target specific spinal cord segments by cross-referencing individual anatomy with a pre-operative CT image. Although the spinous processes serve as an anatomical marker, their narrow and pointed bony structures introduce acoustic challenges, causing irregular reflection, refraction, and scattering of ultrasound waves that contribute to low FUS transmission through the vertebrae^79,126,127^. To overcome this issue while using a single-element transducer, we implemented a hook-and-loop strap system to rotate the marmoset’s torso and spine based on CT image measurements. This directed the ultrasound beam through the vertebral laminae, whose geometry is more amenable to ultrasound transmission^80,126^ (**Figure 6D**). These designs allowed for reliable and spatially precise BSCB disruption across spinal tissue in four marmosets, as detailed below. To further support implementation across the marmoset field, the M-FRAME platform integrates with commercially available hardware including the Rk-50 Marmoset FUS machine (FUS Instruments Incorporated, Toronto, ON, Canada) and the Narishige marmoset stereotax (Model SR-AC, Narishige International Incorporated, Amityville, New York, USA).

## FUS BSCB Disruption

### Identification of Target Coordinates using Pre-FUS CT

Targets for BSCB disruption were selected using a preprocedural CT of the head and neck collected using a Bruker Si-78 small animal PET/CT (Bruker BioSpin Corp, Billerica, MA, USA) equipped with a console running ParaVision 360 (version 3.7, Bruker BioSpin Corp, Billerica, MA, USA). For CT imaging, marmosets were fixed in plane via the imaging component of M-FRAME. CT images were acquired with the following parameters: field of view = 79.6 x 81.1 - 209.3 mm, pixel size = 200 μm, X-ray source filter = 0.5 - 1 mm aluminum, frame averages = 1, scanning mode = step and shoot, rotation angle = 1.0°. Using vertebral landmarks, we selected either single (Marmosets V, H, and M) or multiple (Marmoset S) targets within the cervical and thoracic spinal cord for BSCB disruption. To estimate the rotation required to avoid the spinous processes at each FUS target, we measured the distance from the center of the spinal cord target to the superficial surface of the skin within each animal’s CT. The average soft tissue thickness from the dorsal surface of the skin to the spinous process was ∼1.34 mm and was minimally compressed under a 0.1 kg load (two steps of the motor controlling the transducer position). Once localized on the marmoset with a positioning pointer, target coordinates (x: L/R, y: D/V, z: A/P) were rotated about the spinal cord (z axis) 30° counterclockwise using a matrix calculation (**Figure 6A, B**, **Equation 1**).

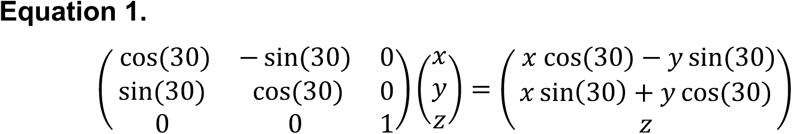

These coordinates were entered into the treatment planning workstation running MORPHEUS software (MORPHEUS framework, FUS Instruments Incorporated, Toronto, ON, Canada) which offered a visual reference to accurately rotate the spine by 30°. To disrupt the BSCB in multiple regions of the spinal cord, translation of the transducer along the rostral-caudal axis was calculated based on the location of preceding points.

### FUS Procedure

To transiently disrupt discrete regions of the BSCB, marmosets were positioned in an RK-50 Marmoset apparatus (FUS Instruments Incorporated, Toronto, ON, Canada) incorporating a marmoset specific stereotax (Model SR-AC; Narishige International Incorporated, Amityville, New York, USA) outfitted with the FUS component of our M-FRAME platform. The FUS apparatus used in these experiments has been described in detail previously^67,88,128^. In brief, an automated 3-axis position system guided a single-element spherically focused ultrasound transducer (500 kHz or 1.5 MHz; FUS Instruments Incorporated, Toronto, ON, Canada) rigidly mounted to the positioning system for precise control of the location of FUS application. The number of pulses, repetition period, and number of bursts commanded by the MORPHEUS software were generated by an external waveform generator (Siglent SDG 1032X, Siglent Technologies, Solon, Ohio, USA) and sent through a 15 W amplifier, matching circuit, and then to the transducer. The transducer was immersed in degassed water via a 300 mL polyimide reservoir which was coupled to the back using degassed, warmed ultrasound transmission gel (01–50, Parker Laboratories, INC., Fairfield, NJ, USA). Water was degassed using a FUS-DS-50 portable water degasser (FUS Instruments Incorporated, Toronto, ON, Canada).

Prior to all FUS procedures, the skin surrounding the target(s) was cleared of hair using a combination of electric clippers and depilatory cream (Nair Shower Cream, Church & Dwight Co, Inc., Ewing, NJ, USA). Immediately prior to FUS application, a bolus of lipid microbubbles (350 - 400 μL/kg; Definity, Lantheus Medical Imaging, Billerica, MA, USA) - prepared by mixing 0.86 mL of sterile saline with 0.1 mL of microbubble stock solution - was administered via the implanted catheter to aid in BSCB disruption. In instances where multiple regions were targeted within the same animal, microbubbles were injected before application of FUS to each target, with 5 minutes between applications. The microbubble solution was cleared from the catheter hub via a 200 μL sterile saline flush. Following microbubble injection, the ultrasound transducer was oriented directly above the appropriate target for immediate application of FUS. BSCB disruption with the 550 kHz transducer employed the following parameters: acoustic pressure = 0.75 - 1.2 MPa, burst duration = 20 ms, burst period = 1000 ms, number of bursts = 70 bursts. BSCB disruption with the 1.5 MHz transducer employed the following parameters: acoustic pressure = 1.8 - 2.5 MPa, burst duration = 20 ms, burst period = 1000 ms, number of bursts = 70 bursts. All reported acoustic pressures are not derated – we have shown that the marmoset skull derates by ∼47% with a 1.5 MHz transducer ^88^.

## Anatomical Imaging of FUS BSCB Disruption

### Post-FUS MRI Acquisition

To visualize BSCB disruption following application of FUS, anesthetized marmosets were transferred from the FUS apparatus to a 9.4 T 30 cm horizontal bore MRI scanner (Bruker BioSpin Corp, Billerica, MA, USA) fit with a Bruker BioSpec Avance Neo console and the software package Paravision-360 (version 3.7; Bruker BioSpin Corp, Billerica, MA, USA). Marmosets were fixed in stereotactic position using the rotatable stereotax included in the M-FRAME platform. After scanner localization, shimming and setup 100-150 μL of a gadolinium-based contrast agent (1 mmol/mL, Gadavist or Gadobuterol, Bayer Healthcare Pharmaceuticals, Leverkusen, Germany) was administered to each animal via the implanted venous catheter. A 200 μL saline flush was used to clear contrast agent from the catheter hub. For imaging, both radiofrequency transmission and reception were accomplished with a two-channel 86 mm inner diameter coil (Bruker BioSpin Corp, Billerica, MA, USA). To detect gadolinium extravasation within the regions of disruption, a magnetization-prepared rapid gradient echo (MPRAGE) pulse sequence was used. Animals receiving FUS BSCB disruption in a single region were imaged with the following parameters: TR = 6,000 ms, TE = 3 ms, field of view = 40 x 33 x 15 mm, matrix size = 160 x 132 x 60 voxels, voxel size = 0.25 x 0.25 x 0.25 mm, bandwidth = 50 kHz, flip angle = 12°, total scan time = 11-minutes 24-seconds. To image multiple BSCB openings in the same animal, a wider field of view was used with the following parameters: TR = 6,000 ms, TE = 3.47 ms, field of view = 50 x 33 x 16 mm, matrix size = 200 x 132 x 64 voxels, voxel size = 0.25 x 0.25 x 0.25 mm, bandwidth = 50 kHz, flip angle = 12°, total scan time = 12-minutes 6-seconds. A minimum of two scans of BSCB disruption were acquired for averaging during processing. These imaging procedures balanced the small size of the marmoset spine with (∼5 mm width at the cervical enlargement, and ∼3 mm in thoracic spine) with imaging acquisition time to limit the spread of gadolinium across the parenchyma^129^ and time spent under anesthesia. To aid in placing regions of BSCB disruption within the internal anatomy of the spinal cord, we have also generated a multimodal marmoset spine template (see **Multimodal Marmoset Spine Template Generation**).

### Anatomical MRI Processing and Analysis

Anatomical scans were first converted from digital images and communications in medicine (DICOM) format to neuroimaging informatics technology (NIfTI) format using the Analysis of Functional NeuroImages (AFNI, version AFNI_23.3.09 ‘Septimius Severus’) software package (dcm2niix_afni, version v1.0.20230411) ^130–132^. All scans were then reoriented using the FMRIB Software Library (FSL, version 6.0.7.18 (modified)) ^133,134^. Anatomical scans were denoised in MATLAB (version 24.2.0.2923080 (R2024b) Update 6). Denoising was based on well validated procedures^66,135,136^ and employs a variance-stabilizing transformation (VST) framework optimized for Rician-distributed noise. ^137^ Briefly, we applied the forwards VST to transform heteroscedastic Rician noise into approximately homoscedastic Gaussian noise. Subsequently, transformed images were denoised via a block-matching and 4D filtering (BM4D) algorithm^138^. To restore the initial intensity range of the denoised images, we then applied the inverse VST.

To acquire measurements of BSCB disruption extent, denoised anatomical scans of the spine were registered to a base image using FSL (FLIRT, version 6.0). After registration, images were averaged using AFNI (3dcalc). Two images were averaged per animal. Average scans were then manually straightened along the rostral-caudal axis using FSL (FSLeyes), and the extent of each locus of BSCB disruption was assessed. The extent of BSCB disruption on *ex vivo* spinal cords was calculated^139^ using coordinate sets from individual animal MRI and CT scans (FSL’s FSLeyes). These measurements were employed for gross anatomical assessment of tissue quality post-FUS (see **Tissue Harvest and Histochemical Staining**).

To compare BSCB disruption at varied ultrasonic pressures and central frequencies, we calculated disruption volume at each treatment location. To do so, we generated a spherical region of interest (ROI) centered within each area of BSCB disruption (FSL’s fslmaths) in preprocessed, post-FUS MR images from Marmosets V and S. Each ROI was of 0.75 mm radius. In each image, we extracted an intensity value from every voxel within each ROI (AFNI’s 3dmaskdump). We then calculated the intensity value corresponding to the lower third quantile for each ROI, which was subsequently used to threshold manually generated masks of BSCB disruption extent at each treatment location (FSL’s fslmaths). Following manual optimization of the resulting mask, the volume of all remaining voxels was calculated using AFNI (3dBrickStat). Quantile calculations were conducted using R (version 4.3.1 (2023-06-16), Beagle Scouts)^140^ in RStudio (version 2024.04.1+748 (2024.04.1+748), Posit Software, PBC) ^141^.

## Systemic AAV Administration

Immediately after verification of BSCB disruption via MRI-based visualization of GBCA extravasation within the targeted site, Marmosets M and H each received a systemic injection of AAV(s) as a bolus through an implanted venous catheter (**Table 1**). Specifically, Marmoset H received a pool containing both AAV9-hSyn-hM3D(Gq)-mCherry (50474, Addgene, Watertown, MA, USA) and AAV9-CAG-GFP (37825, Addgene, Watertown, MA, USA). Marmoset M received just AAV9-hSyn-hM3D(Gq)-mCherry (50474, Addgene, Watertown, MA, USA). For each animal, viral pools were cleared from the catheter hub using a flush of 200 µL sterile saline. At the conclusion of each FUS experiment, the skin was covered with Aquaphor ointment (Beiersdorf Inc., Wilton, CT, USA) to preclude irritation resulting from application of the depilatory cream.

## Marmoset Behavior

### Behavioral Assessment of Upper Limb Motor Function

To assess the effects of FUS application to the spinal cord of Marmoset H, in which the brachial region of the spinal cord was targeted, we conducted a motor assessment of the arm/hand known as the Valley Task^118^. Valley Task testing utilized a dedicated testing apparatus consisting of two components: a modified nest box that is affixed directly to the home cage and a plexiglass Valley Task attachment fit to the nest box^114^. The Valley Task attachment features a single vertical opening through which marmosets may reach rewards at one of five ascending levels on the attachment’s right- and left-hand side. As the vertical opening is centrally placed, marmosets must use the right arm to grasp rewards on the left side of the attachment and *vice versa* for the right arm.

Prior to testing, the order of assessment was determined via random selection for the initial limb after which the side of testing alternated until two runs for the left and right arm were collected for each session. At the beginning of each session, the marmoset was allowed to voluntarily enter the testing apparatus via an opening at the rear of the nest box. Each run began when an opaque plastic divider, which occluded the marmoset’s view of the Valley Task attachment, was removed. The run persisted until the marmoset successfully retrieved each marshmallow reward, positioned centrally at each ascending level of one side of the attachment, or after a maximum of 5-minutes elapsed. Between runs, the plastic divider was replaced, and rewards were positioned at the appropriate side of the attachment according to the alternating pattern described above. At the conclusion of the session, the marmoset was allowed to return to the home cage.

### Behavioral Scoring and Analysis

To assess motor competency of the upper limbs post-FUS, we calculated two metrics from videos collected during completion of the Valley Task (see above): latency and accuracy. Latency was calculated in seconds from the start of the session to the end of the session - the time at which the final marshmallow to be collected arrived at the mouth - or when 5-minutes had expired. In assessing accuracy, the maximum score was 15-points; the final accuracy score was the sum of points earned for collecting each food reward minus the sum of points deducted for errors. One point was awarded for collection of the reward on first level of testing apparatus with points increasing by an increment of one for each successive level to a maximum of five points. Errors decreasing the final score included drops of the food reward after collection and failed grasping maneuvers. Deducted points were only additive if the marmoset removed its hand completely from the apparatus between attempts or if the marmoset, possessing the reward, dropped the reward after a failed grasping maneuver. If the marmoset did not acquire the marshmallow with the proper hand, it received no points.

Statistical analyses of behavioral data were conducted using R in RStudio with the lme4 (version 2.0.6) ^142^ and lmerTest (version 3.1.3) ^143^ packages. Linear mixed-effects models were employed to examine the relationship between fixed effects of arm, time post-FUS (zeroed) and their interaction on Valley Task score and latency. As each session included four trials (two for each arm), each model included session as a random effect. Models were estimated using Restricted Maximum Likelihood. P-values for fixed effects were obtained via an ANOVA and evaluated using an α of 0.05.

## PET Index of DREADD Function

### [^18^F]-FDG PET Image Acquisition

To index the functional effect of actuating transduced chemogenetic receptors (28 days after viral delivery), we conducted FDG PET imaging with Marmoset H after systemic DCZ administration. PET data were collected via a Bruker Si-78 small animal PET/CT (Bruker BioSpin Corp, Billerica, MA, USA) and a console running ParaVision 360 (version 3.7, Bruker BioSpin Corp, Billerica, MA, USA). The animal was fixed within the PET/CT using a custom-built, marmoset-specific stereotax^114^. Immediately prior to PET scanning, the marmoset received a bolus of DCZ (260 µL/kg) followed by a bolus of 17.88 MBq (0.48 mCi) FDG through an implanted venous catheter. PET data were acquired for 90-minutes with a field-of-view of 85 x 85 x 150 mm and were dynamically reconstructed using maximum likelihood-expectation maximization (MLEM 0.5 mm) in bins: 3x20; 4x30; 2x60; 5x300; 6x600 seconds. To localize foci of high FDG uptake within the spine cord in relation to vertebral landmarks, a CT image was collected at the conclusion of PET data acquisition. The CT imaging parameters were as follows: field of view = 79.6 x 81.1 mm, pixel size = 200 μm, X-ray source filter = 0.5 mm aluminum, frame averages = 1, scanning mode = step and shoot, rotation angle = 1°.

### [^18^F]-FDG PET Processing and Analysis

Both PET and CT images were converted from DICOM to NIFTI format and reoriented using a method identical to that for MRI scans. Dynamically reconstructed PET data were then converted to standard uptake values normalized by body weight (SUVbw) and averaged from frames 15-20 (30-minutes to 90-minutes) using AFNI (3dcalc). The post-FUS anatomical MRI, PET, and CT images were then manually cross registered using FSL (FSLeyes) to enable localization of regions of modulated FDG uptake in relation to the position of BSCB disruption. The registered CT image was then used to create a tissue mask of the spinal cord (AFNI’s 3dcalc and FSL’s FSLeyes). Using AFNI, the spinal cord tissue mask was smoothed via sequential dilation and erosion by 1 level (3dmask_tool), before being de-obliqued (3dWarp) and applied (3dcalc) to PET data resampled (3dresample) to CT space (de-obliqued).

To preform statistical analysis on tissue-isolated PET data, we extracted weight normalized FDG uptake (AFNI’s 3dmaskdump) from voxels within 0.5 mm radius spheres placed both within the region of BSCB disruption and at successive vertebral levels along the anterior-posterior extent of the spine (FSL’s fslmaths). Two ROIs, spaced at ∼2 mm, were placed within the region of BSCB disruption. Using R in RStudio, values from ROIs were preprocessed, combined by region (T3 and T2; T1 and C7 (region of BSCB disruption); C6 and C5; C4 and C3) and tested in aggregate for normality using a Shapiro-Wilk test (shapiro_test)^144^. Due to a non-normal data distribution, comparisons of weight-normalized FDG uptake between regions were conducted using a paired-samples Wilcoxon signed rank test with a Bonferroni correction for multiple comparisons (wilcox_test)^144^. For all statistical tests on FDG data, we employed an α of 0.05.

## Tissue Harvest and Histochemical Staining

### Perfusion and Tissue Extraction

At the conclusion of FUS BSCB experiments, Marmosets S and V were euthanized via injection of 390 mg/mL pentobarbital sodium and 50 mg/mL phenytoin sodium (100 mg/kg, EUTHASOL, Virbac, Westlake, TX, USA). Animals receiving systemic AAV injections after FUS BSCB disruption (Marmosets M and H) were euthanized in the same fashion after 4 weeks to allow for robust transgene expression^117^. Immediately following euthanasia, marmosets were perfused transcardially with cold 4% paraformaldehyde (PFA). Each spine was extracted via resection of the vertebral spinous processes at the pedicle and post-fixed for a period of 24-hours in 4% PFA at 4 °C. Post-fixed spinal cords were cryoprotected via submersion in ascending concentrations of sucrose (10% - 30%). Prior to embedding, cryoprotected spinal cords were visually assessed for damage resulting from FUS BSCB disruption. Subsequently, spinal cords were partitioned into sections, embedded in tissue freezing medium and frozen via immersion in dry ice. For tape transfer histochemical staining, spinal cords were embedded in cutting compound (Tissue Plus O.C.T., 23-730-571, Thermo Fisher Scientific Inc., Waltham, MA, USA). For free floating immunofluorescence staining, spinal cords were embedded in clear TFM-C tissue freezing medium (General Data Healthcare, Cincinnati, OH, USA).

### Tape Transfer Histochemical Staining

To preserve the anatomy and order of individual sections of the marmoset spinal cord (Marmosets V and S), we employed a tape transfer assisted cryo-sectioning technique originally developed in rodents^145^ and optimized for the marmoset brain^146^. Briefly, sections were sliced at 50 µm using a stage-modified Leica CM3050S crytostat^146^ located in an environmentally controlled room set to 60% humidity and 18 °C. Throughout the duration of slicing, the interior of the cryostat chamber was set to -22 °C and the spinal cord specimen was maintained at a temperature between -15 and -17 °C. To conduct both Nissl and hematoxylin and eosin (H&E) staining simultaneously within regions of BSCB disruption, we collected three separate series of spinal cord sections (two for staining and one as a contingency) by dividing consecutive slices onto separate slides. Each series contained every third section, resulting in a 150-µm interval between sections on the same slide. Tissue slices were cured to the slide via application of UV light for 12-seconds inside a UV-LED station contained within the cryostat chamber. After curing, slides rested for a period of 24-hours at 4 °C. In total, 627 serial spinal cord sections were collected from Marmoset V (209 per-series) and 1395 sections (465 per-series) were collected from Marmoset S.

To assess damage within regions of BSCB disruptions, we conducted Nissl and H&E staining on separate series of spinal cord tissue. Both Nissl and H&E staining were automated via a Sakura Tissue-Tek Prisma Plus (6130 & Glas g2 6500 Slide Stainer Glass Coverslipper Workstation, Sakura Finetek USA, Inc., Torrance, CA, USA). The Nissl staining protocol employed here has been described previously^145^. In brief, slides were immersed in solution containing 1.88 g thionine acetate (high purity, ThermoScentific Cat. NO. 22973998) dissolved in 750 mL of deionized water, 9 mL of glacial acetic acid (012-00245, FUJIFILM Wako Pure Chemical Corporation, Osaka, Japan), and 1.08 g of sodium hydroxide (221465–500G, Sigma-Aldrich, St. Louis, MO, USA) for a period of 1-minute 25-seconds. Slides were then washed three times in deionized water (45-seconds per-wash), dehydrated in increasing concentrations of ethanol (2-minute washes of 50% and 70%, followed by 2x two-minute washes in 95% and 100%), and cleared via xylene (2x three-minute washes) ^147,148^ before automated cover slipping with a Sakura Tissue-Tek Glas g2 Coverslipper, (Sakura Finetek USA, Inc., Torrance, CA, USA) using Sakura Tissue Tek Mounting Media (Sakura Finetek USA, Inc., Torrance, CA, USA). For H&E staining, we utilized materials from a H&E stain kit (Gil’s II Hematoxylin stain kit, Sakura Finetek USA, Inc., Torrance, CA, USA) and the associated protocol provided by the manufacturer. Briefly, hematoxylin was applied for a period of three-minutes and 15-seconds, after which slides were rinsed in distilled water (2x for one-minute). Then, a weak acid differentiator solution was used for 30-seconds before a bluing reagent was applied to slides for one-minute followed by an additional one-minute rinse in distilled water. Slides were then immersed in 95% alcohol for one-minute prior to application of a highly concentrated eosin Y solution for five-seconds. Finally, slides were rinsed and dehydrated in three changes of absolute alcohol (three-minutes each). The cover slipping procedure was the same as that described for Nissl-stained slides.

### Tissue Sectioning for Fluorescence Microscopy

To verify viral expression within the sites of BSCB disruption, embedded spinal cords from Marmosets M and H were sectioned at 50 µm for free-floating storage in phosphate buffered saline (PBS) using a Leica CM3050 S cryostat (Leica Microsystems, Boston, MA, USA). The cryostat chamber temperature was set to −22 °C, and the object temperature was set to −17 °C. Tissue sectioning was performed in a temperature- and humidity-controlled chamber maintained at approximately 67 °F (19.4 °C) and 60% relative humidity. Free-floating sections both within and outside the region of BSCB disruption were selected for immunofluorescence imaging.

### Microscopy and Computational Reconstruction for Histopathological Assessment

Images of histochemical and immunofluorescence stained sections were captured using a Hamamatsu NanoZoomer S60v2 (C16600-01, Hamamatsu Photonics K.K., Hamamatsu City, Japan). Brightfield imaging was conducted with a 20x objective with a numerical aperture of 0.7, yielding a scan resolution of 0.46 µm/pixel. Images were collected as a stack with 9 focus layers. Fluorescence imaging was conducted using a tissue-appropriate objective and numerical aperture. DAPI, FITC, and TxRed signals were acquired sequentially as separate monochrome images using their respective filter sets. The resolution of fluorescence images was 0.46 µm/pixel. A X-Cite (Excelitas X-Cite 110LED) light source was used to excite each fluorophore.

Histological images were transferred to a data acquisition system (analogous to that described in^146^) for quality assessment and preprocessing as described below. To preprocess each histological series, we employed an automated pipeline originally developed for the marmoset brain^146^. In short, images were cropped to isolate individual tissue sections, converted to tagged image file (TIFF) format, and cross registered (2D alignment) using large deformation diffeomorphic metric mapping (LDDMM) ^149^. An initial rigid-body alignment^150^ was performed to establish the approximate correspondence between adjacent sections, followed by LDDMM-based nonlinear registration. The spatial transformations generated from registration of the down-sampled images were then scaled and applied to the corresponding high-resolution images, allowing the original image data to be transformed into the registered coordinate space without performing computationally intensive registration directly at full resolution. The resulting cross-registered high-resolution serial sections were subsequently assembled to reconstruct a three-dimensional volume of the spinal cord (see **Figure 5B** for an example reconstruction). To isolate stained neural tissue within this reconstructed volume, we applied (AFNI’s 3dcalc) an automatically generated and manually refined (FSLeyes) binary mask. The isolated volume was then reoriented using FSL.

To perform a microscale assessment of the safety of noninvasive FUS BSCB disruption, images of Nissl and H&E stained sections were manually inspected. Nissl-stained sections obtained from regions within and outside the area of blood–spinal cord barrier (BSCB) disruption were systematically examined to assess overall tissue integrity and evidence of cellular injury. Histological evaluation included assessment of tissue architecture, neuronal and cellular morphology, preservation of normal cell boundaries and Nissl substance, and the presence of abnormal nuclear features indicative of cellular damage, including nuclear condensation, shrinkage, fragmentation, or other alterations in nuclear morphology. These observations were used to compare the extent of tissue and cellular damage between regions associated with BSCB disruption and adjacent or remote regions in which BSCB integrity was preserved. H&E sections from the same regions were examined for microhemorrhage as indicated by erythrocyte leakage into the surrounding tissue.

## Multimodal Marmoset Spine Template Generation

### Perfusion, Tissue Extraction and Preparation

To harvest tissue for high-resolution imaging, the template marmoset was euthanized via injection of 390 mg/mL pentobarbital sodium and 50 mg/mL phenytoin sodium (100 mg/kg, EUTHASOL, Virbac, Westlake, TX, USA). Proceeding euthanasia, the marmoset was perfused transcardially via cold 0.1 M phosphate buffer followed by 4% PFA. The spine was then dissected in a continuous tissue block and stored in a custom-designed conical tube containing 4% PFA at 4 °C. After a period of 30-days, the spine block was submerged in a solution of 1X PBS (PBS pH 7.2 (10X), 70013032, Thermo Fisher Scientific Inc., Waltham, MA, USA) and 0.2% GBCA (1 mmol/mL, Gadavist or Gadobuterol, Bayer Healthcare Pharmaceuticals, Leverkusen, Germany) at 4 °C. To allow for adequate absorption of the contrast agent, this solution was replaced after 7-days. Following a period of 12-days, the tissue block was then submerged in a solution of distilled water and 0.2% GBCA to acquire ultra-high resolution anatomical MRI.

### Ex Vivo CT Acquisition of the Marmoset Spine

To acquire *ex vivo* CT of the marmoset spine, the tissue block was placed within a Bruker Si-78 small animal PET/CT (Bruker BioSpin Corp, Billerica, MA, USA) outfitted with a console running ParaVision 360 (version 3.6, Bruker BioSpin Corp, Billerica, MA, USA). Following placement within the PET/CT, both a low- and high-resolution CT image were acquired. The low-resolution CT image was acquired with the following parameters: field of view = 50.2 × 167.6 mm, pixel size = 200 μm, X-ray source filter = no filter, frame averages = 1, scanning mode = step and shoot, rotation angle = 1.0°. The high-resolution CT image was acquired with the following parameters: field of view = 50.2 × 167.6 mm, pixel size = 100 μm, X-ray source filter = no filter, frame averages = 10, scanning mode = step and shoot, rotation angle = 0.8°. Figure 7A shows the template CT by way of 3D-rendering.

**Figure 7.**
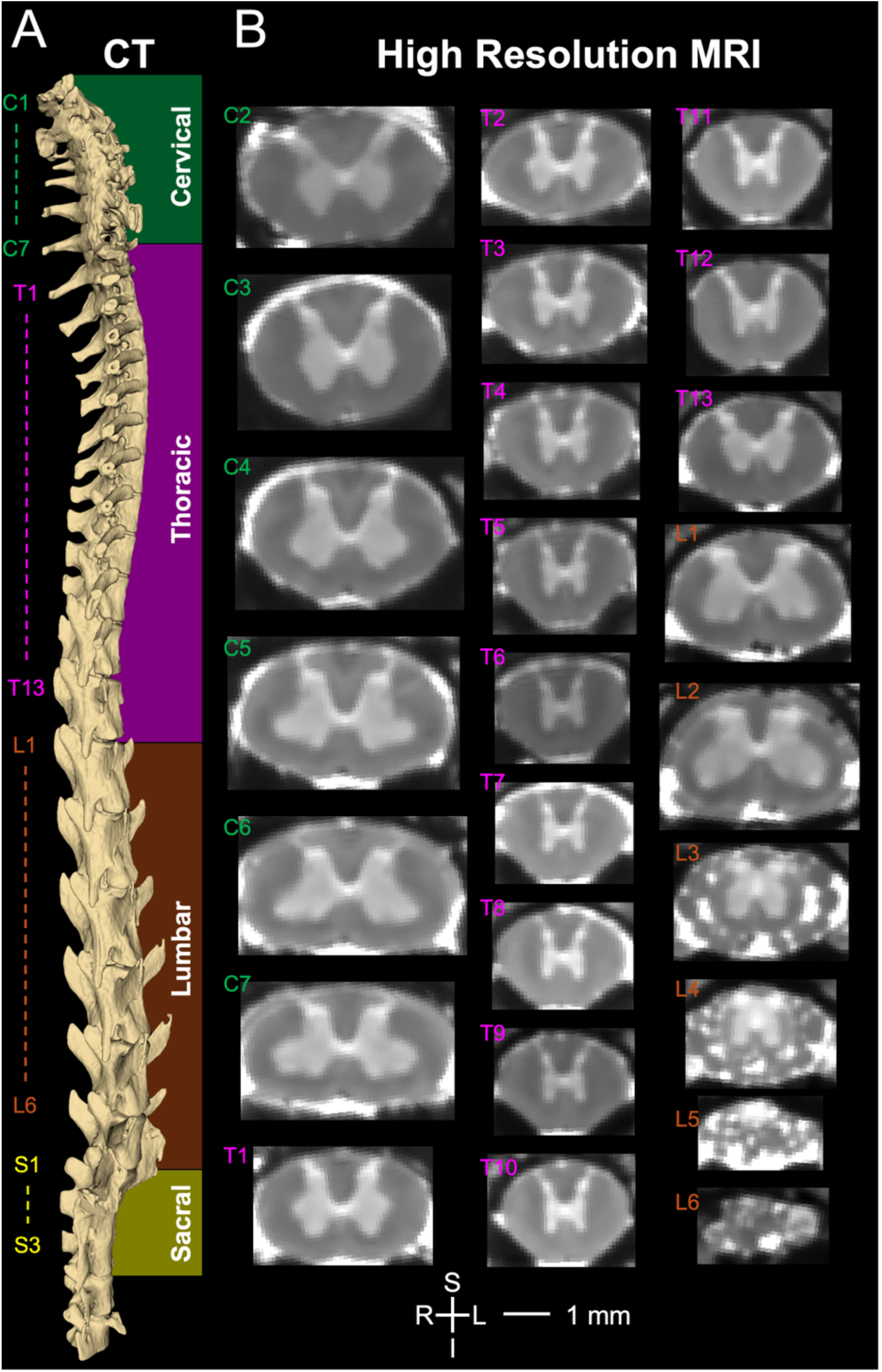
Multimodal template of the marmoset spine. **(A)** 3D-rendering of our high-resolution (100 µm), *ex vivo* CT template of the marmoset spine. The first and last vertebra of the cervical, thoracic, lumbar, and sacral regions of the spine are labeled to the left of the rendering. **(B)** High-resolution (74 µm) anatomical MR images sampled from vertebral levels C2 through L6 of our marmoset spinal cord MRI template. Scale bar = 1 mm for MR images. Abbreviations: S, superior; I, inferior; L, left; R, right; C, cervical; T, thoracic; L, lumbar; S, sacral

### Ex Vivo MRI Acquisition and Processing

To acquire ultra high-resolution *ex vivo* anatomical MRI of the marmoset spine, the tissue block was fixed within a 9.4 T 30 cm horizontal bore MRI scanner (Bruker BioSpin Corp, Billerica, MA, USA), equipped with a Bruker BioSpec Avance Neo console and the software package Paravision-360 (version 3.7; Bruker BioSpin Corp, Billerica, MA, USA). A custom 30 mm inner diameter millipede quadrature coil was used for imaging (ExtendMR LLC, Milpitas, CA, USA). A rapid-acquisition with relaxation enhancement (RARE) pulse sequence was used to image the tissue block with the following parameters: TR = 400 ms, effective TE = 25.53 ms, field of view = 44 x 25.6 x 20 mm, matrix size = 592 x 344 x 270 voxels, voxel size = 74 x 74 x 74 µm, bandwidth = 100 kHz, excitation flip angle = 90.0°, rare factor = 4, total scan time = 5-hours 9-minutes and 36-seconds. Six successive images were acquired to capture the entire length of the spine. *Ex vivo* anatomical scans were pre-processed as outlined above (see ***Anatomical MRI Processing and Analysis***). Following MRI denoising, all scans were smoothed with a 120 µm gaussian kernel in AFNI (3dmerge) for visualization of coronal sections at successive vertebral levels (**Figure 7B**).

## Acknowledgements

We thank Brianne L. Stein, Lauren Dubberley and Dr. Julia Oluoch for animal care and preparation. This work was supported by the National Institute of Neurological Disorders and Stroke of the National Institutes of Health under award number R21NS125372 (D.J.S.) and the National Institute of General Medical Sciences of the National Institutes of Health under award number T32 GM142630 (M.R.C.).

## Author Contributions

M. R. C. Experimentation, Data Analysis, Writing, Editing

S. S. G. Experimentation, Writing, Editing

N. T. N. Experimentation, Writing, Editing

I. Z. R. Experimentation

L. L. Experimentation

D. S. MRI Sequence Development

V. P. C. MRI Denoising

L. Li. Experimentation

M. C. N. Experimentation, Editing

T. K. H. MRI Sequence Development

E. P. Study Conception, Editing

R. S. Study Conception, Editing

M. K. L. Experimentation, Editing

D. J. S. Study Conception, Experimentation, Writing, Editing

