## Supplemental Figure 1 for "Noninvasive Focal Gene Delivery of Functional Neural Actuators to the Primate Spinal Cord using Focused Ultrasound"

**Supplemental Figures**

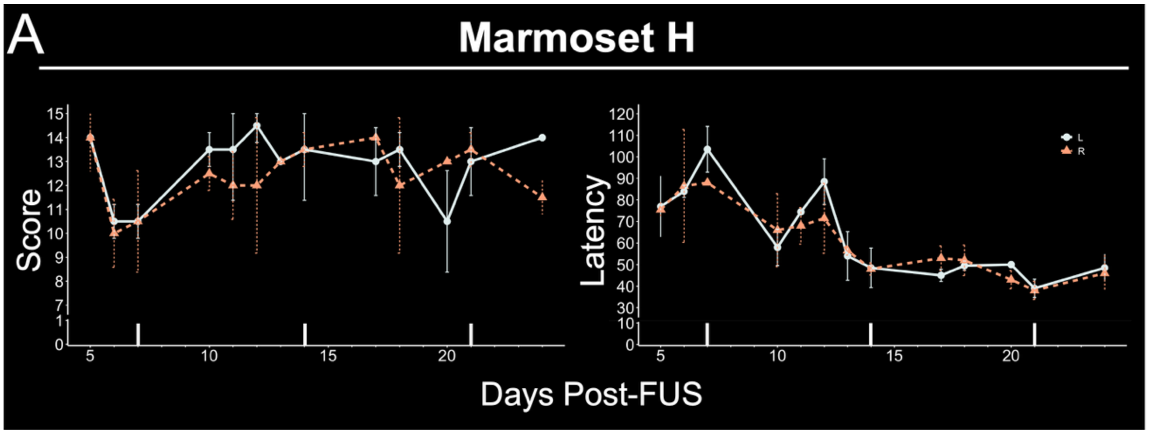


**Supplemental Figure 1. Valley Task score and latency by arm for Marmoset H. (A)** Valley Task[118] score (left) and latency (right) assessed from days five to 24 post-FUS (mean ± SD). Score and latency plots are stratified by arm (left, pale blue; right, orange).
