## Supplemental Table 1 for "Noninvasive Focal Gene Delivery of Functional Neural Actuators to the Primate Spinal Cord using Focused Ultrasound"

### Supplemental Tables

**Supplementary Table 1.** Animal information.

| Marmoset | Sex | Age (months) | Weight (g) |
|---|---|---|---|
| V | M | 27 | 220 |
| S | F | 59 | 205 |
| H | M | 63 | 285 |
| M | F | 40 | 235 |
| Template | F | 52 | 230 |
